# Filopodia powered by class X myosin promote fusion of mammalian myoblasts

**DOI:** 10.1101/2021.07.28.454163

**Authors:** David W. Hammers, Cora C. Hart, Michael K. Matheny, Ernest G. Heimsath, Young il Lee, John A. Hammer, Richard E. Cheney, H. Lee Sweeney

**Author notes:** Corresponding Author Correspondence: H. Lee Sweeney, 1200 Newell Dr. ARB R5-216, Gainesville, FL 32610-0267, Phone: -392-3558.

## Abstract

Skeletal muscle fibers are multinucleated cellular giants formed by the fusion of mononuclear myoblasts. Several molecules involved in myoblast fusion have been discovered, and finger-like projections coincident with myoblast fusion have also been implicated in the fusion process. The role of these cellular projections in muscle cell fusion was investigated herein. We demonstrate that these projections are filopodia generated by class X myosin (Myo10), an unconventional myosin motor protein specialized for filopodia. We further show that Myo10 is highly expressed by differentiating myoblasts, and Myo10 ablation inhibits both filopodia formation and myoblast fusion *in vitro*. *In vivo*, Myo10 labels regenerating muscle fibers associated with Duchenne muscular dystrophy and acute muscle injury. Conditional loss of *Myo10* from muscle-resident stem cells, known as satellite cells, severely impairs postnatal muscle regeneration. Furthermore, the muscle fusion proteins Myomaker and Myomixer are detected in myoblast filopodia. These data demonstrate that Myo10-driven filopodia facilitate multi-nucleated mammalian muscle formation.

## INTRODUCTION

The development, growth, and repair of vertebrate skeletal muscle is largely mediated by the ability of myoblasts to fuse with each other and with pre-existing muscle fibers. In postnatal muscle, this fusion process is initiated by the activation of muscle-resident stem cells, known as satellite cells, that normally remain in a quiescent state positioned between the sarcolemma and basement membrane of muscle fibers (Mauro, 1961). Following an activation stimulus, satellite cells give rise to myoblast progeny which proliferate, differentiate, and fuse to bring the muscle back to homeostasis (Charge and Rudnicki, 2004).

The fusion of mammalian myoblasts requires the merging of two apposing lipid bilayers and has been shown to involve several widely-expressed protein classes, including cytoskeleton elements (Charrasse et al., 2006; Randrianarison-Huetz et al., 2018; Vasyutina et al., 2009), phagocytosis receptors (Hamoud et al., 2014; Hochreiter-Hufford et al., 2013; Park et al., 2016), and calcium-sensing membrane repair proteins (Leikina et al., 2013; Posey et al., 2011). Myomaker (Millay et al., 2013) and Myomixer [aka Myomerger and Minion; the product of the *Gm7325* gene (Bi et al., 2017; Quinn et al., 2017; Zhang et al., 2017)] have been identified as being membrane proteins essential for myoblast fusion. Current evidence suggests Myomaker is required by both fusing cells, while the requirement for Myomixer is only unilateral (Quinn et al., 2017). However, the mechanisms regulating the function of these proteins are currently unknown. Thin, actin-filled projections have been observed during the fusion of murine (Randrianarison-Huetz et al., 2018) and zebrafish (Gurevich et al., 2016) myoblasts. These structures appear to be filopodia, which are thin, membrane enclosed projections of actin bundles that are important for cellular behaviors such as path finding during migration, interaction with the extracellular matrix, and cell to cell communication (Mattila and Lappalainen, 2008). The actin cytoskeleton is well-established as an enactor of *Drosophila* myoblast fusion (Abmayr and Pavlath, 2012; Chen, 2011), with filopodia suggested to be involved in this process (Segal et al., 2016). Despite actin cytoskeletal involvement in the fusion of both vertebrate and arthropod myoblasts, no members of the myosin superfamily of molecular motors have been reported to have a direct role in the muscle fusion process.

Myosin superfamily members perform actin-associated functions in all cell types. This includes the conventional (Class II) myosins, such as those that power muscle contraction, and several classes of “unconventional” myosins, of which eleven are expressed in mammals (Berg et al., 2001; Odronitz and Kollmar, 2007). These unconventional myosins employ the same basic motor mechanism as conventional myosins, but have unique tail domains that allow them to perform specialized cellular functions. Class X myosin (Myo10) is an unconventional myosin involved in the formation and elongation of filopodia in mammalian cells (Berg and Cheney, 2002). Myo10 consists of a N-terminal motor domain, a lever arm with three calmodulin binding sites and a single alpha helical domain, and a C-terminal tail containing a pleckstrin homology (PH), a myosin tail homology 4 (MyTH4), and a 4.1/Ezrin/Radixin/Moesin (FERM) domain (Kerber and Cheney, 2011). Upon activation, Myo10 forms an anti-parallel dimer that is optimized for the organization and movement along actin bundles (Ropars et al., 2016). Myo10-driven filopodia are involved in processes such as neural and vascular development (Heimsath et al., 2017; Pi et al., 2007; Zhu et al., 2007).

Because filopodia-like structures have been observed in fusing muscles (Gurevich et al., 2016; Randrianarison-Huetz et al., 2018), we sought to determine if these structures are, indeed, Myo10-driven filopodia involved in skeletal muscle fusion. In this work, we show that Myo10, a filopodia associated myosin, is a key component of myoblast fusion. Myo10 deficiency in myoblasts results in loss of detectable filopodia and muscle fusion *in vitro* and impaired muscle regeneration *in vivo*. Lastly, we demonstrate that the fusion proteins Myomaker and Myomixer can be detected within muscle filopodia.

## RESULTS

### Protrusions from differentiating myoblasts are apparent during cellular fusion

Thin actin-filled cellular extensions that protrude from differentiating myoblasts have been observed during vertebrate myofusion (Randrianarison-Huetz et al., 2018). We sought to investigate the occurrence and behaviors of these projections in living myoblasts of both undifferentiated and differentiated states via live-cell confocal microscopy, utilizing membrane targeted fluorescent reporter constructs containing a C-terminal human H-Ras CAAX box prenylation signal (RFP-CAAX or GFP-CAAX; see Methods) to enable detailed visualization of cellular protrusions. Undifferentiated myoblasts of the murine C2C12 cell line exhibit many thin cellular projections. Live imaging reveals that these projections are actively elongating primarily at the leading edge of myoblasts, while the trailing regions of the cells predominantly exhibit retraction fibers that become evident as the cells move (**Figure S1A-C**, **Supplemental Movies 1-2**; summarized in **Figure 1A**), as commonly observed during cell migration (Mattila and Lappalainen, 2008). Upon induction of differentiation by switching the cells to low-serum media conditions, myoblasts undergo morphological changes characterized by cellular elongation, loss of distinct directional polarity, and increased incidence of cellular projections from the entirety of the cell body (**Supplemental Movie 3**). As the differentiation process proceeds to the formation of multi-nucleated myotubes, the cells display an array of dynamic and static projections at the lateral edges, prominent dorsal protrusions along the cell body, and arm-like lamellipodial extensions adorned with thin projections that can protrude from any part of the cell (**Figure 1B** and **Supplemental Movies 4-7**; Summarized in **Figure 1C**). The lengths of extending projections from myotubes are significantly longer than those from the leading edge of myoblasts (**Figure S1D**), and scanning electron micrographs confirm that these dorsal projections from myotubes are of consistent structure and size of thin cellular projections known as filopodia (**Figure 1D**). Live imaging of differentiating myoblast cultures reveals the involvement of these structures in muscle cell fusion, as witnessed through fusion induced by projections extending from the lateral edge of myotubes (**Figure 1E-F**, **Supplemental Movie 8**), as well as by projection-laden lamellipodial extensions (**Figure 1G, Supplemental Movies 9-10**). This evidence demonstrates that these cellular projections extending from differentiating myoblasts are involved in the formation of multi-nucleated mammalian muscle. The remainder of this report will focus on the cellular mechanisms responsible for the generation of these projections and their role in muscle fusion.

**Figure 1.**
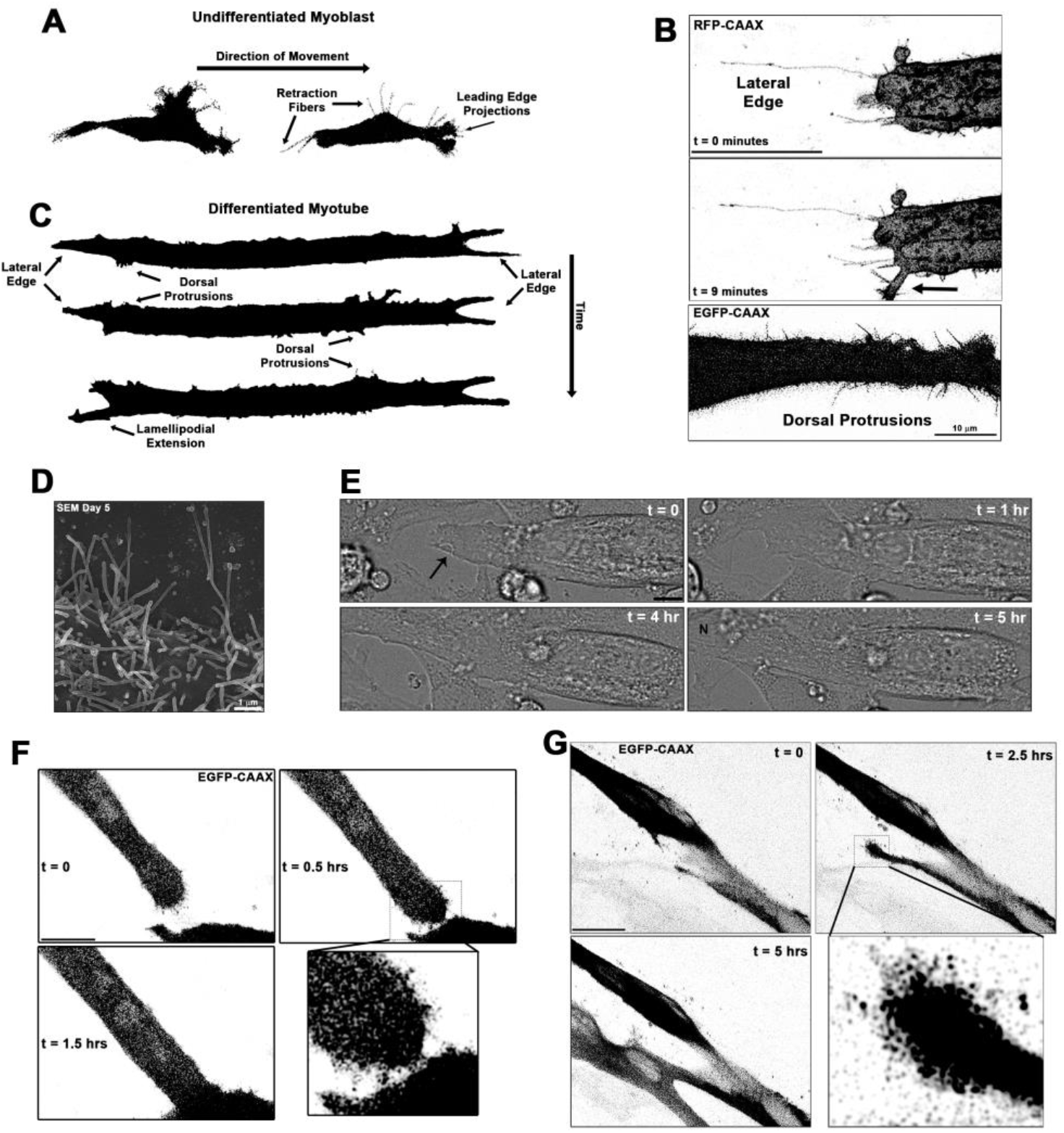
Cellular projections are prominent on differentiating muscle cells and participate in cell fusion. (**A**) A summary schematic depicting the cellular protrusions exhibited by undifferentiated myoblasts. (**B**) Live-cell confocal microscopy of differentiated myotubes (Day 5) expressing a membrane-targeted fluorescent protein constructs (RFP-CAAX or GFP-CAAX) reveals myogenic projections are dynamic structures featured across the cell surface, including prominent lateral edge projections, dorsal protrusions, and those emerging from lamellipodial extensions (indicated by arrow). (**C**) Summary schematic displaying cellular projections associated with differentiated myotubes. (**D**) Cellular projections visualized on the surface of differentiating myoblasts by scanning electron microscopy (SEM). (**E**) Differential interference contrast imaging of a myotube exhibiting lateral edge protrusions (indicated by arrow) actively engaged in myoblast fusion (N indicates newly incorporated nucleus; Day 4-5). Fluorescently-labeled myoblasts utilizing (**F**) lateral edge protrusions and (**G)** a lamellipodial extension adorned with fine protrusions to promote fusion with adjacent cells (differentiation Days 4-5). EGFP-CAAX in (**G**) becomes transferred to the non-expressing cell upon fusion, making the newly added cell visible via fluorescence. Unless otherwise noted, scale bars represent 25 µm.

### Myo10 is required for filopodia formation and cellular fusion of myoblast in vitro

A hallmark of filopodia is the presence of Myo10, which is a molecular motor associated with the initiation and elongation of filopodia and potential cargo binding within filopodia (Berg and Cheney, 2002; Zhang et al., 2004). To establish if the projections we visualize during myogenic fusion are, indeed, filopodia, we investigated the expression pattern of Myo10 within differentiating myoblast cultures. Myo10 protein content (**Figures 2A**, **S2A**) and *Myo10* gene expression (**Figure 2B**) increase during the time course of myoblast differentiation. Myo10 immunofluorescence localizes specifically to differentiated myotubes (**Figure 2C**), which are confirmed to have expression of myosin heavy chain (MHC; **Figure 2D**). The Myo10-positive projections observed on these myotubes also contain actin filaments (F-actin; **Figure 2E**), a key feature of filopodia.

**Figure 2.**
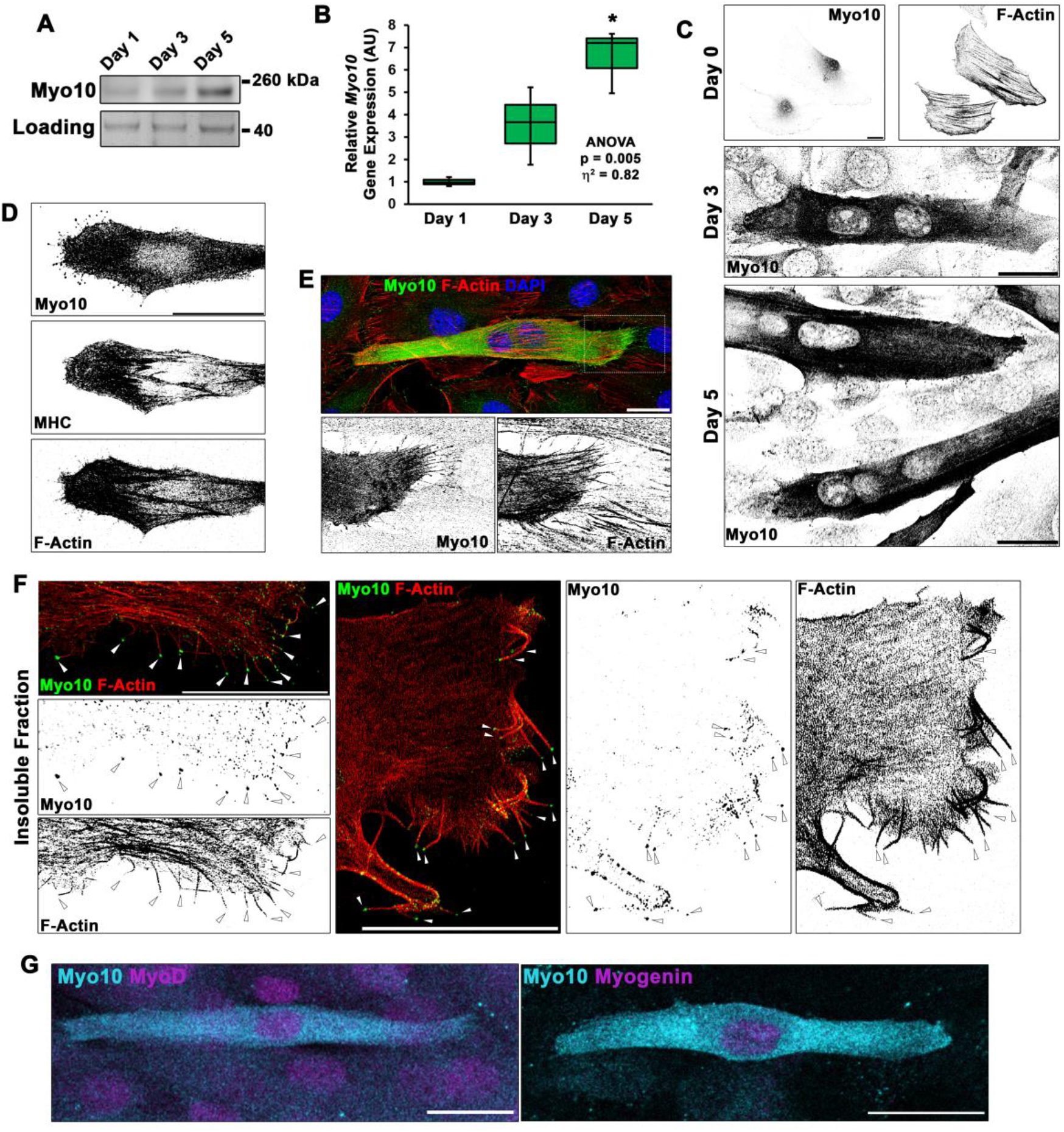
Differentiating myoblast cultures express Myo10. (**A**) Myo10 protein content, as shown by immunoblotting, and (**B**) *Myo10* gene expression, as measured by real-time PCR (n = 3 independent cultures for each time point; normalized to *Gapdh*), is increased during the myoblast differentiation time course. (**C**) Immunofluorescence (IF) reveals that the increase in Myo10 in differentiating myoblast cultures is localized primarily to differentiated myotubes, which (**D**) also express the muscle terminal differentiation marker, myosin heavy chain (MHC). (**E**) The Myo10-positive cellular extensions of differentiated myoblasts contain F-actin. (**F**) IF of the insoluble fraction of differentiating myoblasts reveals that insoluble Myo10 is found distinctly at the tips of F-actin bundles (indicated by arrows). (**G**) Myo10-positive myoblasts exhibit nuclear staining for both MyoD and Myogenin muscle regulatory factors, as shown by IF, following one day of exposure to differentiation conditions. Gene expression data are presented as box-and- whisker plots depicting 2^nd^ and 3^rd^ quartiles with minimum and maximum values (relative to Day 1 values). Data were analyzed using one-way ANOVA followed by Tukey post-hoc tests [α = 0.05; *p <0.05 vs Day 1 values; effect size is presented as eta-squared (η^2^)]. Unless otherwise noted, scale bars represent 25 µm.

Evidence suggests Myo10 can exist as an inactive, diffusible folded monomer that undergoes a conformational change during activation that allows for unfolding and anti-parallel dimer formation, resulting in engagement with the actin cytoskeleton (Ropars et al., 2016; Umeki et al., 2011). Because Myo10 appears to fill the entire cell of differentiated myotubes, we sought to determine if myocyte Myo10 represents a freely-diffusible population, an actin-bound population, or combination of the two states. Fractionation of differentiating myoblast cultures into soluble and insoluble cellular fractions revealed that Myo10 partitions into both the soluble and insoluble fractions (**Figure S2B**), with a slightly larger proportion residing in the soluble fraction. Serving as fractionation controls, αTubulin partitions primarily into the soluble cellular fraction, while MHC and actin are predominantly found in the insoluble fraction (**Figure S2B**). Immunofluorescence of insoluble myotube cellular remnants following soluble fraction extraction reveals that insoluble Myo10 is associated with the actin cytoskeleton (**Figure S2C**), and can be distinctly visualized at the tips of actin bundles that appear to be within myotube filopodia (**Figure 2F**).

In agreement with Myo10 expression becoming highly activated in myoblasts during myogenic differentiation, analysis of the full-length *Myo10* promoter region (Lai et al., 2013) revealed the presence of 14 consensus E-Box motifs (CANNTG; depicted in **Figure S2D**). These motifs are DNA elements bound by myogenic regulator factors (MRFs), such as MyoD and Myogenin, during myogenesis (Tapscott, 2005). Co-expression of constitutive GFP-CAAX with a mApple (RFP) construct driven by the *Myo10* promoter was used to investigate Myo10 activation in myoblasts exposed to differentiation medium for one day compared to those undergoing differentiation for four days. Activation of the *Myo10* promoter, as determined by the RFP/GFP-CAAX ratio in immunoblots, is confirmed to increase proportionally with Myo10 content as myoblast differentiation progresses (**Figure S2E-F**). Live cell imaging early in the differentiation time course (Day 1) revealed individual myoblasts with low basal expression of mApple detaching from the culture substrate, undergoing a transition into a blebbing spherical morphology with increased mApple expression, and re-attachment to the culture substrate in a morphology resembling differentiated myocytes (**Figure S2G**, **Supplemental Movie 11**). Consistent with MRF-mediated activation of Myo10 during myoblast differentiation, Myo10-positive mononuclear myocytes exhibit positive staining for both MyoD and Myogenin, which have both been shown to bind to the *Myo10* promoter (Cao et al., 2006), during the first day of differentiation (**Figure 2G**). These data indicate that *Myo10* is activated early in the differentiation period of myogenesis.

The requirement of Myo10 for the formation of muscle filopodia was investigated in *Myo10* knockdown experiments using C2C12 myoblast cell lines generated by lentiviral-mediated expression of control or *Myo10-*targeted short-hairpin RNA (shRNA) and clonal selection. *Myo10* knockdown (KD) results in efficient loss of *Myo10* gene expression during both growth (**Figure S2H**) and differentiation (**Figure S2I**) culture conditions, as well as reduction of Myo10 protein from both culture conditions (**Figure 3A**) and loss of Myo10 immunofluorescence during differentiation (**Figure 3B**). Myogenic differentiation potential *per se* is not affected by loss of Myo10, MHC protein and *Myh2* gene expression do not differ between control and *Myo10* KD lines after 5 days of differentiation (**Figure 3A**, **S2I**). Loss of Myo10 does, however, have a significant effect on prevalence of cellular projections, which are therefore filopodia, as *Myo10* KD myocytes exhibit less dorsal protrusions (**Figure 3C**) and no detectable extending filopodia (**Figure 3D**; **Supplemental Movie 12**). The primary cellular protrusions displayed by *Myo10* KD cells appear as blebs reaching to the cell periphery in order to create connections to the surface substrate (**Supplemental Movie 13**), which are significantly shorter in length when compared to the thin and distinct Myo10-driven filopodia of control shRNA myocytes (**Figure 3D-E**). Loss of Myo10 does not prevent the formation of lamellipodial extensions; however, they are devoid of detectable thin projections, which are thus confirmed to be filopodia (**Supplemental Movie 13**).

**Figure 3.**
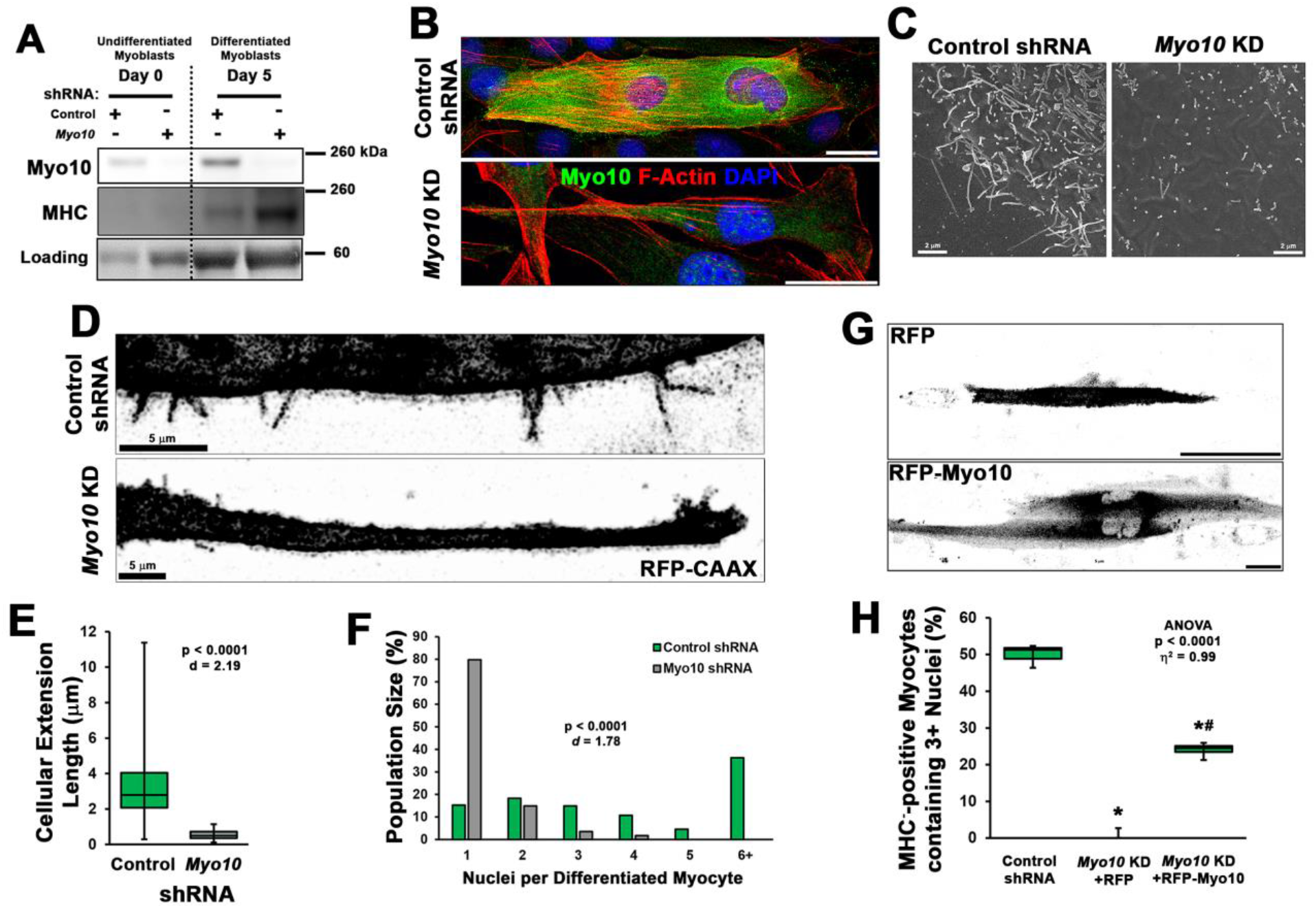
Loss of myoblast Myo10 prevents filopodia formation and cellular fusion. Clonal lines of C2C12 cells expressing control or *Myo10*-targeted shRNA were validated for efficacy of *Myo10* knockdown (KD) and myogenic differentiation potential. (**A**) Immunoblotting for Myo10 protein and the myogenic differentiation marker, myosin heavy chain (MHC; loading control visualized by Ponceau Red staining). KD of *Myo10* myoblasts results in loss of filopodia during differentiation compared to control shRNA cells, as demonstrated by (**B**) immunofluorescence, (**C**) scanning electron microscopy (n = 31-152 cellular extensions), and (**D**) live-cell confocal microscopy, as well as loss of (**E**) cellular extension lengths. Myoblast differentiation assays (n = 3 individual experiments) reveal loss of multinucleated myotubes formation in *Myo10* KD cells after 7 days of differentiation compared to control cells, quantified as (**F**) population distribution of myotube nuclear content. (**G**-**H**) Loss of fusion ability by *Myo10* KD cells can be partially rescued by transfection of a full-length Myo10 construct with an N-terminal mApple fluorescent tag (RFP- Myo10; n = 3-6 individual experiments). Data analysis performed using (**E-F**) Welch’s 2-tailed T- test (α = 0.05) with effect size displayed as Cohen’s d (*d*) or (**H**) one-way ANOVA followed by Tukey post-hoc tests [α = 0.05; *p <0.05 vs control values; ^#^p < 0.05 vs. RFP values; effect size is presented as eta-squared (η^2^)]. Unless otherwise noted, scale bars represent 25 µm.

The involvement of filopodia in myoblast fusion is also confirmed, as loss of Myo10 nearly abolishes multinucleated myotube formation following seven days of differentiation (**Figure 3F**). This can be partially rescued by expression of exogenous Myo10, using an N-terminal mApple tagged human Myo10 construct (RFP-Myo10; **Figure 3G-H**, **S2J**), as determined by the quantification of MHC-positive myocytes containing three or more nuclei following seven days of differentiation. We chose this threshold of myonuclei content as an indication of fusion since prior studies have reported the presence of bi-nucleated myocytes following differentiation of myoblasts lacking the fusion proteins, Myomaker (Millay et al., 2013) or Myomixer (Bi et al., 2017). The rescue of *Myo10* KD myoblast fusion by exogenous Myo10 expression is not attributed to solely filopodia formation, but requires Myo10’s cargo binding functions, as a truncated RFP- Myo10 construct lacking the C-terminal PH, MyTH4, and FERM domains (RFP-Myo10ΔCBD) does not restore myoblast fusion despite induction of filopodia (**Figure S3**). Furthermore, the requirement of myoblast Myo10 for fusogenic activity appears to be a requirement of both fusing cells, as a mixture of control and *Myo10* KD myoblasts having distinct fluorescent labels rarely results in the fusion of the two populations (**Figure S4**). Together, these data reveal that the unconventional myosin, Myo10, is important for muscle formation *in vitro*.

### Myo10 labels regenerating muscle fibers in vivo

Given the robust impact of Myo10 loss on myoblast fusion, we next investigated Myo10- dependent skeletal muscle processes *in vivo*. Myo10 expression in postnatal regenerative myogenesis was examined in the muscle sections from the *mdx* mouse, a mouse model of Duchenne muscular dystrophy (DMD) that continuously displays regions of stable, damaged, and regenerating muscle fibers within the same muscle section, due to loss of dystrophin (Hoffman et al., 1987; Petrof et al., 1993). Strong Myo10 immunoreactivity within small muscle fibers of regenerating areas, which are identified by the presence of centrally-located nuclei (Coulton et al., 1988), occurs in these muscle sections, with negligible signal detection in regions of stable muscle fibers (**Figure 4A**). This Myo10 expression pattern is also observed in human muscle, as DMD patient biopsy samples contain many small, Myo10-positive fibers localized to regenerating foci (**Figure 4B**). Thus, Myo10 is expressed in muscle during times that are expected to have high amounts of myoblast fusion, including muscle regeneration.

**Figure 4.**
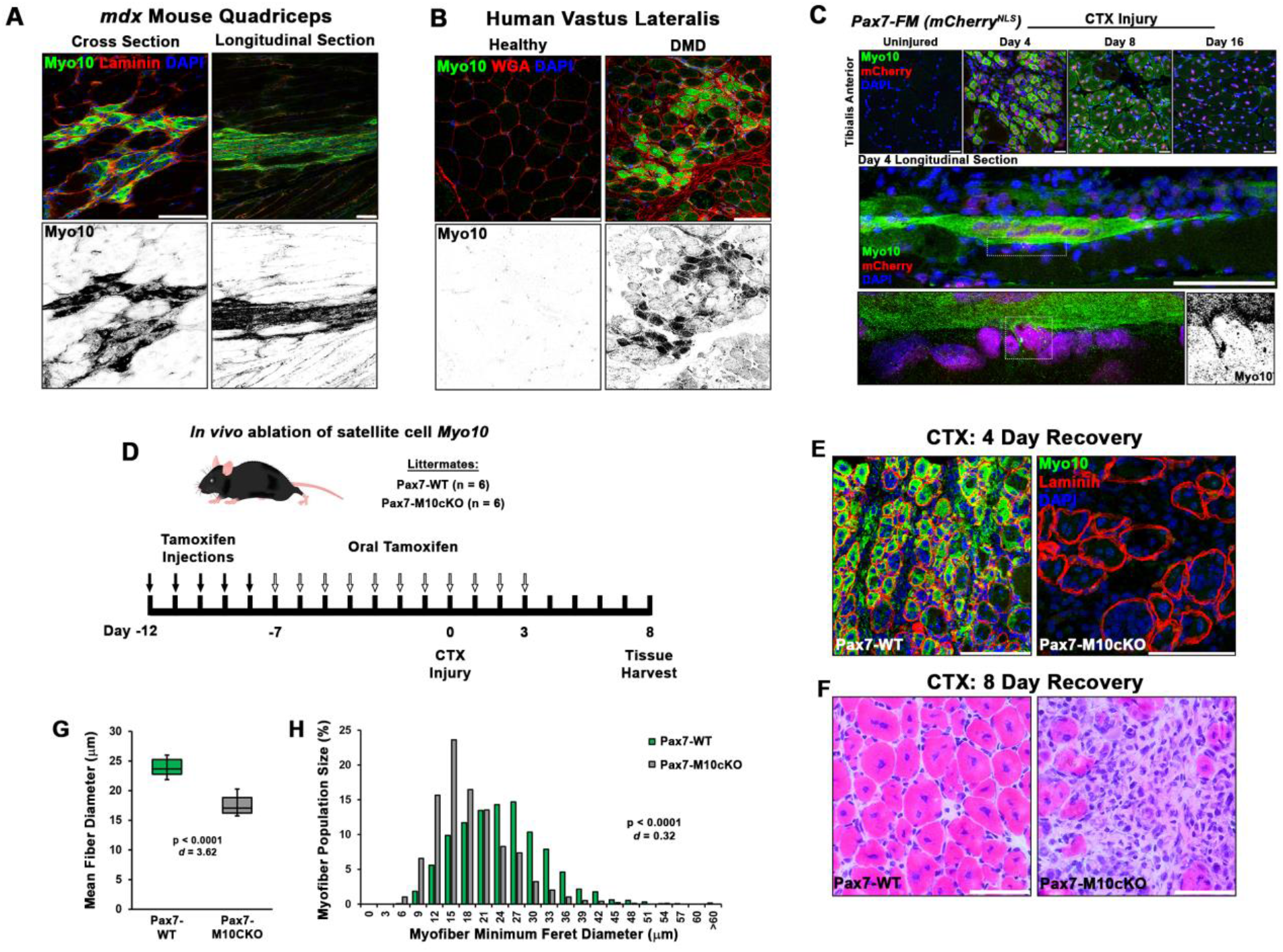
Myo10 labels regenerating muscle fibers *in vivo* and is required for efficient muscle regeneration from injury. Myo10 immunoreactivity in regions of dystrophin-deficient skeletal muscle samples from (**A**) *mdx* mice and (**B**) Duchenne muscular dystrophy (DMD) patients that are undergoing active regeneration. (**C**) Fate-mapping (FM) of muscle satellite cells using the Pax7^Cre-ERT2^ allele crossed onto mice harboring a floxed nuclear-localized (NLS) mCherry allele demonstrates Myo10 expression is found in regenerating muscle fibers of satellite cell origin. The inset shows Myo10-filled filopodia can found extending towards mononuclear myoblasts. (**D**) The role of Myo10 in postnatal muscle regeneration was investigated using Pax7^Cre-ERT2^ conditional *Myo10* knockout (Pax7-M10cKO; n = 6) mice and their non-floxed littermates (Pax7-WT; n = 6). Tamoxifen induction was achieved via 5 consecutive daily intraperitoneal injections of 100 mg/kg tamoxifen (denoted by solid arrows) followed by daily oral treatments with 10 mg/kg tamoxifen (empty arrows) for 7 days preceding cardiotoxin (CTX) injury of the tibialis anterior muscle (TA) and continuing until 3 days after injury. (**E**) This protocol that results in efficient elimination of Myo10^+^ myocytes as evidenced in 4 day recovery muscle. Following 8 days of recovery, Pax7-M10cKO mice demonstrate impaired muscle regeneration compared to Pax7-WT muscle, as evidenced by (**F**) impaired histological recovery and (**G-H**) reduced muscle fiber size (n = 991-1307 fibers). Data are presented as (**G**) box-and-whisker plots depicting 2^nd^ and 3^rd^ quartiles with minimum and maximum values or (**H**) a histogram of entire data set populations, and are analyzed using two-tailed Welch’s T-tests with effect size presented as Cohen’s *d* (*d*). Scale bars represent 100 µm.

Since asynchronous bouts of degeneration and regeneration in parallel characterize dystrophic muscle diseases, an acute model of synchronized muscle damage and subsequent regeneration was employed in non-dystrophic mice to verify that Myo10 of muscle fibers is associated with regenerative myogenesis rather than damage or degeneration. Intramuscular injection of the myotoxin, cardiotoxin (CTX), into the tibialis anterior (TA) of *Pax7^Cre-ERT2^* mice crossed with a nuclear-localized mCherry reporter (Pax7-mCherry^NLS^), a model which allows for satellite cell fate-mapping, was performed. This enables distinct labeling of regenerative myogenesis by identifying mCherry-positive nuclei after tamoxifen-induced Cre-recombinase activation (Nishijo et al., 2009). Post-injury muscle development reveals highly elevated Myo10 levels within regenerating, mCherry-positive myofibers four days following CTX injection (**Figure 4C**). Analysis of muscle after eight and 16 days of regeneration shows Myo10 content to progressively decline back to uninjured levels as the myofibers reach post-regenerative maturation [confirmed via immunoblotting (**Figure S5A**)]. This phenomenon is not exclusive to the CTX model, as intramuscular glycerol injection, an alternative muscle regeneration model, also results in mCherry-positive regenerating myofibers strongly labeled by Myo10 (**Figure S5B**). Therefore, regenerative myogenesis exhibits muscle-specific expression of Myo10 similar to findings *in vitro*. These data also demonstrate that Myo10 is an effective marker to label regenerating skeletal muscle fibers *in vivo*, which may be a useful tool to identify newly formed muscle fibers *in lieu* of developmental MHC isoforms.

### Loss of Myo10 in satellite cells impairs muscle regeneration

The consequence of myoblast-specific loss of Myo10 on muscle regeneration *in vivo* was assessed using *Pax7^Cre-ERT2^* mice crossed to the floxed *Myo10* (*Myo10^tm1cltm1c^*) allele (Heimsath et al., 2017), generating a mouse line capable of inducible ablation of *Myo10* in satellite cells and, thus, their myoblast progeny and any resulting myofibers. In the absence of tamoxifen-induced *Myo10* ablation, homozygous *Myo10^tm1c/tm1c^* mice (termed Pax7-M10cKO for Pax7-<u>M</u>yo<u>10 c</u>onditional <u>k</u>nock<u>o</u>ut) have indistinguishable phenotypes from their *Myo10^tm1c/+^* or *Myo10^+/+^* littermates (termed Pax7-WT) following CTX-induced regeneration in the TA (**Figure S5C-D**). Tamoxifen-induced Cre expression (via the protocol depicted in **Figure 4D**) results in efficient ablation of Myo10 positive muscle fibers (**Figure 4E**) and Myo10 protein content (**Figure S5E**) in Pax7-M10cKO mice four days following CTX injection. Remnants of extracellular matrix from pre-existing muscle fibers, known as ‘ghost fibers’ (Webster et al., 2016), predominate Pax7-M10cKO muscle sections at this time point. At eight days following CTX injection, Pax7-WT muscles demonstrate robust regeneration, while Pax7-M10cKO muscles exhibit impaired regeneration, as evidenced by fewer and smaller myocytes present in regenerating musculature (**Figure 4F-H**). Pax7-M10cKO muscle regenerative defects are also exhibited following freeze injury, a more severe muscle injury model (Hardy et al., 2016), of which affected Myo10-deficient musculature is largely replaced by intramuscular fibrosis following 21 days of recovery (**Figure S6**). Thus, Myo10 is important for regenerative myogenesis.

### Muscle fusion proteins localize to filopodia

A possible role for My10-driven filopodia in muscle fusion is that they provide a means to create cellular contacts required for delivery of the fusion proteins, Myomaker and Myomixer, to apposing cellular membranes at a distance, thus increasing the probability of a fusion event occurring. To investigate this possibility, the localization of these fusion proteins on differentiating myoblasts was assessed via immunofluorescence utilizing commercial antibodies for Myomixer (extracellular epitope; applied prior to cellular permeabilization) and Myomaker (intracellular epitope; applied following permeabilization). The ability of these antibodies to provide specific signals for their respective target proteins was evaluated using exogenous expression of wild-type versions of Myomaker or Myomixer in undifferentiated myoblasts (**Figure S7A-B**).

Mononuclear myocytes early in the differentiation process exhibit strong extracellular staining of Myomixer on most of the cell periphery, including cellular projections (**Figure S7C**). In these cells, Myomaker is localized primarily to vesicular structures that are particularly evident in the perinuclear cap region (**Figure S7C**). This agrees with the previously-reported Golgi localization of Myomaker (Gamage et al., 2017). In multinucleated myotubes, Myomixer remains prominently localized at the cellular periphery, while Myomaker is additionally observed in puncta found in close proximity to the cell membrane along the cell body and cellular projections, including filopodia protruding from lamellipodial extensions (**Figures 5A, S7D**). Myo10-positive cells co-expressing Myomaker and Myomixer are observed during the first day of differentiation (**Figure S7E**), indicating temporal regulation of these myogenic genes are synchronized. The expression of Myomixer and Myomaker are not, however, dependent on Myo10, as both proteins are found in differentiated Myo10 KD myoblasts (**Figure S7F**). In fully differentiated Myo10- positive myotubes, puncta of both Myomaker and Myomixer are observed in Myo10-filled filopodia (**Figures 5B, S7G**). Furthermore, these proteins are found to co-localize with Myo10 puncta in the filopodia remnants of insoluble myotube fractions (**Figure 5C-D**). The localization of both Myomaker and Myomixer to Myo10-positive filopodia is also observed when functional Flag-tagged versions of these proteins, namely the Myomaker-F203 (Millay et al., 2013) and Myomixer- Flag (Zhang et al., 2017) constructs, are exogenously-expressed in differentiating myoblast cultures (**Figure S8**).

**Figure 5.**
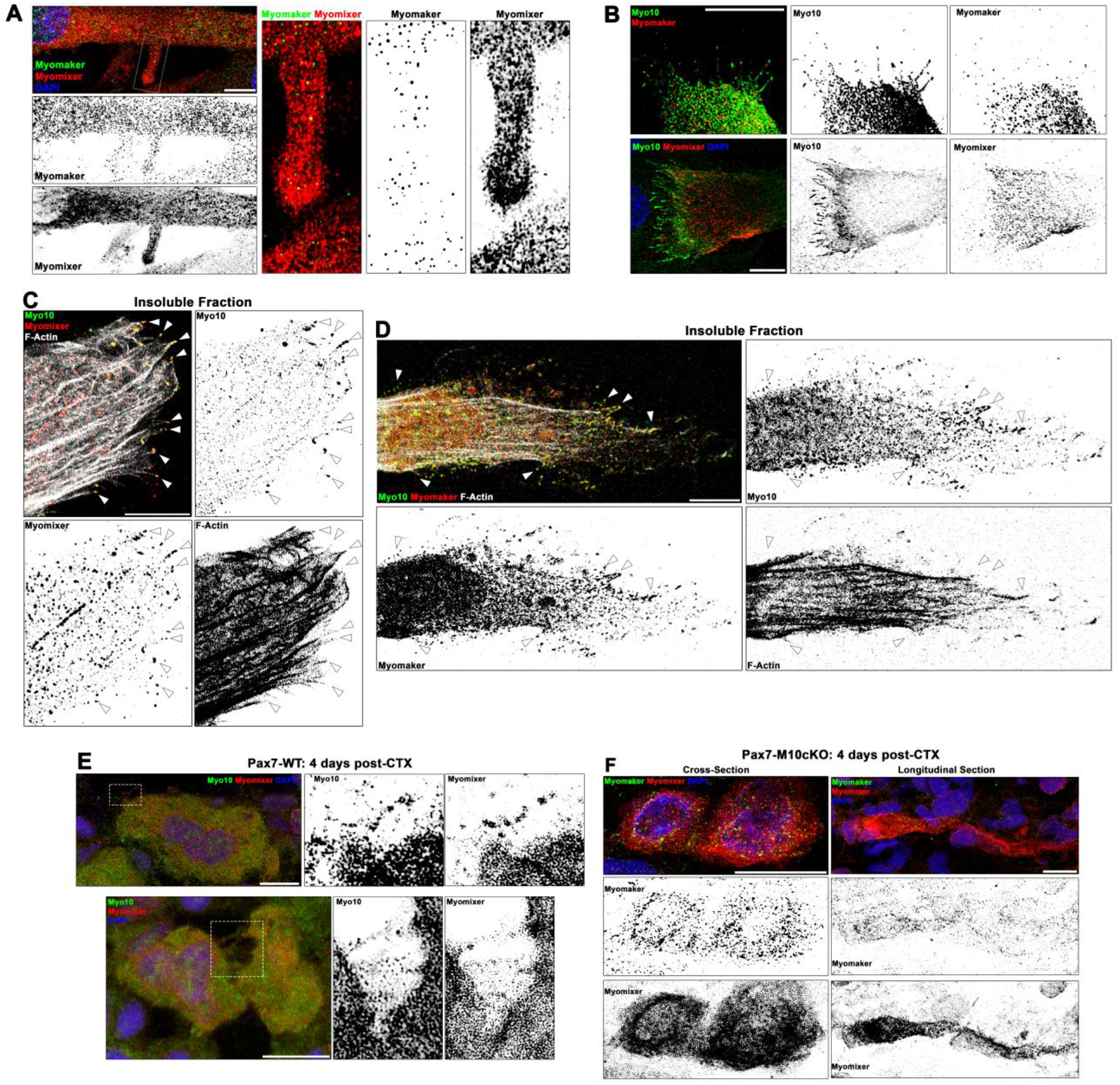
Myogenic fusion proteins are detected along filopodia in differentiating muscle cells. (**A**) Immunofluorescent detection of Myomaker and Myomixer in differentiating myoblast cultures. Inset shows Myomixer and Myomaker puncta on the surface of a lamellipodial extension with filopodia. (**B**) Myomaker and Myomixer puncta are found localized to Myo10-filled filopodia of differentiating myoblasts. Co-localization of (**C**) Myomixer and (**D**) Myomaker with Myo10 in filopodia remnants of the differentiating myoblast insoluble fraction (indicated with arrows). (**E**) Tibialis anterior (TA) muscle cross-sections of Pax7-WT mice at 4 days following cardiotoxin (CTX)-induced injury. (**F**) Cross-section (left) and longitudinal section (right) of Pax7-M10cKO TA muscle at 4 days following CTX-induced injury. Scale bars represent 10 µm.

Sections from regenerating Pax7-WT muscle also display Myo10-labeled filopodia decorated with Myomixer puncta (**Figure 5E**), suggesting this relationship also exists *in vivo*. Such cellular projections are not found on Myomaker- and Myomixer-positive cells of regenerating Pax7-M10cKO muscle (**Figure 5F**), which exhibit thin morphologies similar to *Myo10* KD myoblasts rather than robust myotube formation found in wild-type muscle sections (**Figure 4C**). These findings are consistent with the hypothesis that muscle filopodia provide a means to enable cellular connections that facilitate the function of these fusion proteins upon delivery to a cellular target.

## DISCUSSION

Cellular projections provide cells the ability to explore surrounding space and present molecules for intercellular communication and interaction during tissue development and regeneration. The current work demonstrates that the finger-like projections observed on differentiating mammalian myoblasts are Myo10-driven filopodia that participate in myoblast fusion. Data also demonstrate that myoblast expression of Myo10 is required for formation of multinucleated myotubes *in vitro* and efficient regeneration of skeletal muscle from injury *in vivo*. Furthermore, evidence is provided demonstrating that the fusogenic proteins Myomaker and Myomixer are observed on Myo10-powered filopodia.

Myosin motors compose a diverse protein superfamily with many classes that have all evolved to perform specialized cellular processes. With roles encompassing myocyte contractility, cellular movement, vesicle transport, endocytosis, organelle positioning, and formation and maintenance of filopodia, stereocilia, and microvilli (Sweeney and Holzbaur, 2018), the myosins expressed in humans are essential for many physiological activities. This is highlighted by a large number of diseases resulting from myosin mutations, including cardiomyopathy (Geisterfer-Lowrance et al., 1990; Mohiddin et al., 2004), skeletal myopathy (Armel and Leinwand, 2009), deafness (Friedman et al., 1999; Mohiddin et al., 2004), aneurysms (Pannu et al., 2007), and enteropathy (Golachowska et al., 2012).

While the involvement of the actin cytoskeleton is well documented in both arthropod and mammalian myoblast fusion (Chen, 2011; Millay et al., 2013; Peckham, 2008; Randrianarison-Huetz et al., 2018; Segal et al., 2016), no motor proteins have been previously described to have a direct role in the fusion process itself. Only the conventional myosin, non-muscle myosin II, has been implicated in myoblast fusion, serving as a sub-sarcolemmal mechanosensor (Kim et al., 2015). The current report also provides the first described role of Myo10 in mammalian muscle, which has only been previously detected in muscle via dystrophic muscle gene expression arrays (Marotta et al., 2009) and *Myo10* promoter binding by the muscle regulatory factors MyoD and Myogenin (Cao et al., 2006).

Interestingly, a myosin motor involved in myoblast fusion has not been identified in *Drosophila*, which lack class X myosin. It is likely that another member of MyTH4-FERM containing myosins, such as the class XV myosin (Myo15) homolog *Sisyphus* (Liu et al., 2008), is adapted for the role of filopodia formation in insect muscle. A recent study has demonstrated that loss of Myo15 in *Drosophila* substantially reduces larva viability and causes abnormal neuromuscular junction formation, whereas muscle-specific Myo15 overexpression causes the development of peculiar F-actin structures in myocytes (Rich et al., 2021). While this report did not address whether these actin-based structures are associated with cellular protrusions, the images provided do resemble thin actin bundles reminiscent of myotube dorsal filopodia described in the current study. While the definitive identification of a myosin motor that drives insect muscle filopodia awaits further investigation, the premise of filopodia-facilitated myoblast fusion appears to be a convergent evolutionary feature of multinucleated muscle formation (Segal et al., 2016). Thus the physical presentation of molecules at a distance from the cell body via cytoskeletal extensions is an important aspect of skeletal muscle development as it is in the development of other tissue types (Pi et al., 2007; Zhu et al., 2007).

A paramount finding of this report is the defective muscle regeneration caused by loss of Myo10 in satellite cells. These data are potentially of clinical significance, as a patient having two alleles for a truncating *MYO10* mutation has been recently identified (Patel et al., 2018). While this individual was found on basis of having microphthalmia, a developmental eye defect, it is possible that an underlying myopathy may exist or develop as a result of impairments in muscle recovery from injury. In fact, microphthalmia is also associated with several syndromes that also present with myopathy or neuromuscular disorders, including Walker-Walburg syndrome (Vajsar and Schachter, 2006) and Charcot-Marie-Tooth disease (Fernandez-Torre et al., 2001). Furthermore, several *MYO10* single-nucleotide polymorphisms have been identified (Burghardt et al., 2010), including missense mutations in the N-terminal motor domain. Mutations such as these have the potential to create dominant-negative Myo10 molecules if motor function is impaired or destroyed, and individuals harboring such polymorphisms may exhibit muscle regenerative defects, as demonstrated by loss of *Myo10* in mice. The identification of a Myo10- deficient patient also confirms findings in mice that loss of Myo10 is not absolutely lethal, although less than half of Myo10-null mice survive birth, and those that do survive exhibit developmental deficits (Heimsath et al., 2017). These evidences of decreased Myo10-null embryonic survivability further indicate that Myo10-driven filopodia serve to increase the probability of proper cellular connectivity during developmental processes. The fact that the surviving Myo10-null animals have fused muscle fibers likely indicates that myoblasts are in close opposition during development, lessening the dependence on distal interactions mediated by filopodia. In contrast, satellite cell-derived myoblasts of injured adult muscles must traverse much larger distances in order to locate and fuse with each other, creating a greater dependence on filopodia for postnatal muscle repair than for embryonic muscle development.

Following confirmation that Myo10-driven filopodia are involved in the fusion events required to form multinucleated muscle, we investigated if there is a relationship between these fine cellular projections and the recently discovered muscle fusion proteins, Myomaker and Myomixer. Our experiments reveal that Myomaker and Myomixer are both highly expressed in Myo10-positive myocytes and present in muscle filopodia. It is currently hypothesized that Myomaker’s role in cellular fusion is the promotion of outer membrane leaflet mixing (i.e. hemifusion) between cells, while Myomixer acts as an inducer of intercellular pore formation by promoting positive spontaneous membrane curvature within hemifusion structures (Golani et al., 2021; Leikina et al., 2018). These actions are consistent with the concept of their delivery by Myo10-driven filopodia in order to create an intermediate structure that is essentially a tunneling nanotube, whether configured as individual or bundled lumens (Sartori-Rupp et al., 2019), that precedes development of a full syncytium. Such a structural organization is suggested by the images provided in **Figure 1F**, and Myo10-dependent formation of tunneling nanotubes has been suggested as a requirement for osteoclast differentiation into multinucleated cells (Tasca et al., 2017). While it remains to be confirmed if Myo10 plays an active role in the localization and/or activity of these fusogenic proteins or whether Myo10-driven filopodia increase the probability proper cellular connectivity via increased surface area, the failure of RFP-Myo10ΔCBD to restore myoblast fusogenic activity, despite promoting filopodia formation, suggests Myo10 does have an active cargo-binding role in promoting fusion of mammalian myoblasts.

The findings detailed in this report describe a role of filopodia driven by the unconventional myosin, Myo10, in the formation of multinucleated skeletal muscle. These data further emphasize the importance of these fine cellular projections in the intricate biological processes required for proper development of higher-order organisms, and that perturbations to their formation and function have the ability to cause muscle pathology and potentially modify the course of muscle disease.

## METHODS AND MATERIALS

### Animals

All animal procedures were approved and conducted in accordance to the University of Florida IACUC. C57BL/10 (RRID: IMSR_JAX:000476), *mdx* (RRID: IMSR_JAX:001801), and H2B-mCherry^fl/fl^ (RRID: IMSR_JAX:023139) mice used for this study were from colonies originally derived from Jackson Laboratories. *Pax7^Cre-ERT2^* mice were a generous gift from Dr. Charles Keller (Nishijo et al., 2009). The *Myo10^tm1c^* floxed allele was generated as previously described (Heimsath et al., 2017). Tamoxifen-induced Cre expression for fate-mapping experiments was achieved by intraperitoneal injections of 20 mg/mL tamoxifen (Sigma-Aldrich No. T5648) dissolved in sterilized sunflower seed oil (Sigma-Aldrich No. S5007) at a dose of 100 mg/kg for 5 consecutive days, which results in ∼85-90% labeling efficiency in Pax7 cells. To achieve near 100% induction for conditional ablation studies, mice were subjected to the 5 day injection protocol described above followed by daily oral administration of 10 mg/kg tamoxifen (sunflower seed oil vehicle) starting at seven days preceding injury to 3 days following injury (depicted in **Figure 4D**). Oil only injections and oral treatments served as sham induction controls. Injections and treatments were performed within two hours of the start of the mouse dark cycle to facilitate drug distribution. Only male mice were used for these experiments. The genotypes of all mice used for this study were verified by PCR-based genotyping. Mice were randomly assigned into experimental groups prior to experiments.

Injury of the tibialis anterior (TA) muscle was performed by injecting 50 µL of sterile solutions of either 12 µM cardiotoxin (Calbiochem No. 217503; dissolved in sterile PBS) or 50% glycerol longitudinally through the length of the muscle. Freeze injury was performed by applying a liquid N2-cooled metal rod to the mid-belly of a surgically-exposed TA from a randomly-selected hind-limb for 10 seconds. Following the allocated recovery time from injury, mice were euthanized via CO2. TA muscles were dissected free, either snap-frozen in liquid N2 or embedded in OCT compound and frozen in melting isopentane, and stored at -80 C until analysis.

### Cell culture

C2C12 murine myoblasts (ATCC No. CRL-1772; RRID: CVCL_0188) were purchased from ATCC and used between passages 5 and 13. Cells were cultured at 37 C in 5% CO2 in growth media consisting of high-glucose DMEM (Gibco No. 10566), 10% fetal bovine serum (FBS; Sigma-Aldrich No. F8067), and 1% penicillin/streptamycin (P/S; Gibco No. 15140). C2C12 differentiation media consisted of low-glucose DMEM (Gibco No. 11885), 2% horse serum (Hyclone No. SH30074), and 1% P/S and was changed every two days during differentiation experiments. Ectopic expression experiments were performed using X-tremeGENE 9 DNA transfection reagent (Sigma-Aldrich No. 6365779001) or electroporation using the 4D-Nucleofector system (Lonza). For live-cell imaging, cells were plated on collagen- or gelatin-coated glass-bottom dishes (Willco Wells No. GWST-3522) prior to transfection, differentiated in phenol red-free media (using glutamate-supplemented Gibco No. 11054 DMEM in place of No. 11885), and mounted in a stage-top incubator (Tokai HIT No. INUB-GSI2-F1) with 5% CO2 for image acquisition.

Control or *Myo10* shRNA knockdown C2C12 lines were made using MISSION shRNA lentiviral particles (Sigma-Aldrich No. SHCLNV; Control shRNA ID- SHC002; *Myo10* shRNA IDs- TRCN0000110606 and TRCN0000375033; 5 MOI) following the manufacturer’s directions. Puromycin-resistant clones were selected and verified for *Myo10* knockdown efficiency and myogenic differentiation capacity. For all assays comparing control and *Myo10* KD lines, equal cell numbers were plated and switched to differentiation medium 12-16 hours after plating.

### Plasmids

The tagRFPt-HRAS-CAAX (RFP-CAAX) plasmid was generated by adding the C-terminal 20 amino acids of human H-Ras (NCBI Accession No. NP_005334) to a tagRFP-C vector (Evrogen) modified with a S158T point mutation to enhance photostability (Shaner et al., 2008). EGFP-HRAS-CAAX (GFP-CAAX) was constructed similarly by placing the HRAS-CAAX box motif on the C-terminus of EGFP in the pEGP-C1 vector (Clontech). The RFP-Myo10 construct was prepared by cloning mApple (from Addgene No. 54631) to the N-terminus of human Myo10 (NCBI Accession No. NP_036466) using a G-G-R linker, similar to as previously described (Ropars et al., 2016), in pCDNA3.1(+) vector (Thermofisher No. V79020). The RFP-Myo10ΔCBD construct was prepared by fusing mApple to the N-terminus of a human Myo10 construct lacking the PEST, PH, MyTH4, and FERM domains (aa 1-938), as previously described (Ropars et al., 2016), in pCDNA3.1(+) vector. The *Myo10* reporter plasmid was constructed by replacing CMV promoter of pCDNA3.1(+) with the -1835/+314 region of the *Myo10* promoter region (Lai et al., 2013) followed by an mApple open-reading frame. Murine Myomaker (NCBI Accession No. NP_079652), Myomixer (NCBI Accession No. NP_001170939), Myomaker-F203 (Millay et al., 2013), and Myomixer-Flag (Zhang et al., 2017) open reading frames were cloned into pCDNA3.1(+) vector. All constructs were verified by sequencing and restriction analysis, and all plasmids were prepared in endotoxin-free conditions.

### Immunofluorescence and fluorescent labeling

Tissue immunofluorescence (IF) was performed as previously described (Hammers et al., 2016). Briefly, OCT-embedded frozen muscle was sectioned into either cross-sections or longitudinal sections of 10 µm thickness, fixed in ice-cold acetone, blocked in 5% BSA-PBS + 0.1% Triton X- 100, and incubated in primary antibody overnight at 4 C. Secondary antibodies were applied for 1 hour at room-temperature the following day. Lipofuscin-induced autofluorescence was eliminated using 0.1% Sudan Black B dissolved in 70% ethanol, ensued by a wash in 0.1% Triton X-100 in PBS. Sections were mounted in Vectashield (+DAPI; Vector Labs No. H1200), cover- slipped, and sealed. Control and DMD patient samples were acquired from the National Disease Research Interchange (NDRI; Philadelphia, PA).

Cells cultured on gelatin or collagen-coated coverslips were rinsed twice with PBS, fixed in 4% PFA-PBS for 30 minutes at room temperature, permeablized with 0.1% Triton X-100 in 4% PFA-PBS, blocked with 0.5% BSA-PBS, and incubated in primary antibody overnight at 4 C.

External epitopes of Myomixer were specifically stained by incubation of appropriate primary antibodies for 1 hour prior to permeabilization step. Secondary antibodies were applied for 1 hour at room temperature the following day, followed by 20 minute incubation with Alexa 647- conjugated phalloidin (1:300 in PBS; Life Technologies No. A22287), when appropriate. Cells were counterstained with DAPI and mounted onto pre-cleaned glass slides with Prolong Gold mounting media (Life Technologies No. P36934). Insoluble myotube fractions were prepared for immunofluorescence by removing the soluble cellular fraction via incubation of cells with ice-cold PBS containing 1% Triton X-100, 5 mM EDTA, and protease and phosphatase inhibitor cocktails for 1 hour. The remaining insoluble fraction of the cells was fixed in 4% PFA following careful removal of the soluble fraction and two washes with ice-cold PBS.

Primary antibodies used for IF include anti-Myo10 (0.3 µg/mL; Sigma No. HPA024223; RRID: AB_1854248), anti-Myo10 (2 µg/mL; Santa Cruz Biotechnology No. sc166720; RRID: AB_2148054), anti-Laminin (1.25 µg/mL; Acris Antibodies No. BM6064P), anti-MHC (0.25 µg/mL; R&D Systems No. MAB4470; RRID: AB_1293549), anti-mCherry (1:2000; Novus No. NBP2- 25158; RRID: AB_2636881), anti-MyoD (5 µg/mL; Thermofisher No. MA1-41017; RRID: AB_2282434), anti-Myogenin (2 µg/mL; Novus No. NB100-56510; RRID: AB_838604), anti- TMEM8C/Myomaker (1 µg/mL; Thermofisher No. PA5-63180; RRID: AB_2648742), anti- Myomixer/Myomerger/ESGP (0.67 µg/mL; R&D No. AF4580; RRID: AB_952042), and anti-Flag (1.6 µg/mL; Sigma-Aldrich No. F3165; RRID: AB_259529). Secondary antibodies (all 1:500 dilution) used include Alexa 488 donkey anti-rabbit IgG (Life Technologies No. A21206), Alexa 647 donkey anti-rabbit IgG (Life Technologies No. A31573), Alexa 568 goat anti-mouse IgG (Life Technologies No. A11031), Alexa 568 donkey anti-sheep IgG (Life Technologies No. A21099), and TRITC donkey anti-chicken IgY (Jackson No. 703-025-155). Appropriate primary antibody isotype controls were used in combination of secondary antibodies to ensure specificity of signal. All images were acquired with a Leica SP8 confocal microscope and processed with Leica LAS X software. Image acquisition was performed in sequential scan mode to ensure fidelity of fluorescent signal observed. Cellular projection lengths were analyzed from time-lapse images, where each projection was measured at its longest observed length using FIJI image analysis software (NIH). Image-based quantifications were performed by investigators blind to experimental groups.

### Immunoblotting

Preparation of muscle protein homogenates was performed as previously described (Hammers et al., 2017). Cell lysates for direct immunoblotting experiments were prepared by lysis of cell cultures with SDS-supplemented T-Per lysis reagent (Thermofisher No. 78510) containing protease and phosphatase inhibitor cocktails. For cell fractionation experiments, the soluble cellular fraction was obtained by incubation of cells with ice-cold PBS containing 1% Triton X-100, 5 mM EDTA, and protease and phosphatase inhibitor cocktails. Following removal of this soluble fraction, the insoluble fraction was solubilized using an equal volume of SDS-supplemented T-Per lysis reagent containing protease and phosphatase inhibitor cocktails.

All samples were prepared for SDS-PAGE by boiling in Laemeli’s sample buffer containing 50 mM DTT, run on 4-12% Tris-glycine SDS gels, and transferred to nitrocellulose membranes, as previously described (Hammers et al., 2017). Following blocking in 5% BSA-TBST, membranes were incubated with anti-Myo10 (0.15 µg/mL; No. AF4580; RRID: AB_952042), anti-MHC (0.125 µg/mL; R&D Systems No. MAB4470; RRID: AB_1293549), anti-RFP (0.5 µg/mL; Abcam No. ab62341; RRID: AB_945213), or anti-GFP (2.5 µg/mL; Abcam No. ab13970; RRID: AB_300798) primary antibody overnight at 4° C. Membranes were incubated with HRP-conjugated anti-rabbit IgG (1:1000; Cell Signaling No. 7074), anti-mouse IgG (1:1000; Cell Signaling No. 7076), or anti-chicken IgY (1:1000; Jackson Labs No. 303-035-003) secondary antibody for 1 hour, and developed with ECL reagent (Thermofisher No. 34577). Images were captured using the C-Digit Imaging System (Licor). Ponceau-red staining was used to verify equal loading of comparative samples.

### Real-time PCR

Real-time PCR was performed as previously described (Hammers et al., 2017) using the following mouse-specific primers: *Myo10* (forward) 5’-TTC CAC CGC ACA TCT TCG CCA TTG-3’ and (reverse) 5’-CCC CGG GAT TCT GCC TCA CTA CTC-3’; *Myh2* (forward) 5’-AGA ACA TGG AGC AGA CCG TG-3’ and (reverse) 5’-TCA TTC CAC AGC ATC GGG AC-3’; *Gapdh* (forward) 5’-AGC AGG CAT CTG AGG GCC CA-3’ and (reverse) 5’-TGT TGG GGG CCG AGT TGG GA-3’. Relative gene expression quantification was performed using the ΔΔCt method with *Gapdh* as the reference gene.

### Tissue Histology

OCT-embedded frozen muscle was cross-sectioned into 10 µm thick sections, stained with Hematoxylin & Eosin (H&E) or picrosirius red (PSR) as previously described (Hammers et al., 2020), and visualized with a Leica DMR bright-field microscope equipped with a digital camera (Leica No. DFC480). Image analysis was performed using FIJI software.

### Statistical analysis

Quantified data of this study are displayed as box-and-whisker plots (depicting 2^nd^ and 3^rd^ quartiles with minimum and maximum values) or as histograms of full population distribution, and were analyzed using two-tailed Welch’s T-test (α = 0.05; effect size reported as Cohen’s *d*) or one-way ANOVA (effect size reported as η^2^) followed by Tukey post-hoc tests (α = 0.05). Power analyses (power = 0.8; α = 0.05) using previous or preliminary data for each measure dictated all sample sizes utilized in this study. No data points were excluded from data analysis during the course of this study.

## ACKNOWLEDGEMENTS

This work was funded by R01-AR075637 from the National Institute of Arthritis and Musculoskeletal and Skin Diseases (NIAMS), a Wellstone Muscular Dystrophy Cooperative Center grant (U54-AR-052646) from the National Institutes of Health, and Fondation Leducq funding (13CVD04) to HLS. DWH was supported by the Muscular Dystrophy Association (MDA549004) during the course of this work. REC was supported by a National Institutes of Health grant (NIGMS R01 GM134531), EGH was supported by NCI T32CA009156 to the Lineberger Comprehensive Cancer Center, and JAH was supported by the intramural program of the National Heart, Lung, and Blood Institute. The authors thank Dr. Melissa Merscham-Banda, Radhika Bhake, Lillian Wright, and Dr. Laurence Prunetti for technical assistance during this work.

## AUTHOR CONTRIBUTIONS

DWH, REC, and HLS designed the experiments. DWH, CCH, MKM, YL, and EGH performed the experiments. DWH and HLS performed data analysis. DWH, JAH, REC, and HLS interpreted the data. DWH and HLS drafted the manuscript with contributions from all authors.

## SUPPLEMENTAL FIGURES

**Figure S1.**
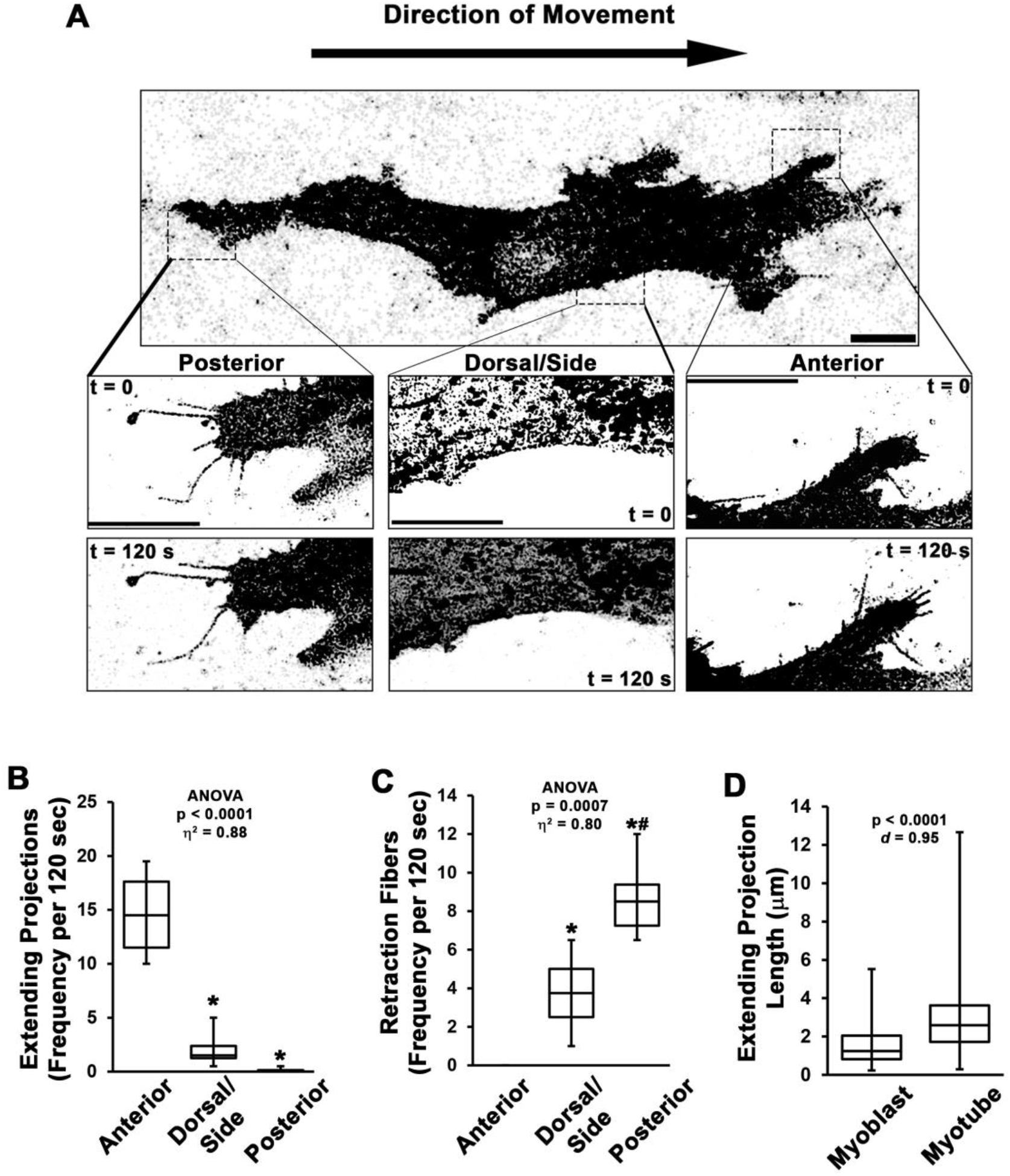
Dynamics of myoblast cellular projections. (**A-C**) Analysis of cellular projections in undifferentiated myoblasts expressing RFP-CAAX reveals elongating cellular projections are primarily observed on the anterior leading edge of the cell, whereas retraction fibers are the predominant cellular extensions observed away from the cell anterior (n = 4 independent experiments). (**D**) Quantification of extending projection length in undifferentiated myoblasts and differentiated myotubes (n = 115-244 projections from 3 individual cultures). Data are presented as box-and-whisker plots depicting 2^nd^ and 3^rd^ quartiles with minimum and maximum values and were analyzed using (**B-C**) one-way ANOVA followed by Tukey post-hoc tests (α = 0.05; *p < 0.05 vs. Anterior values; #p < 0.05 vs. Dorsal/Side values) or (**D**) two-tailed Welch’s T-tests with effect size presented as Cohen’s *d* (*d*). Scale bars represent 5 µm.

**Figure S2.**
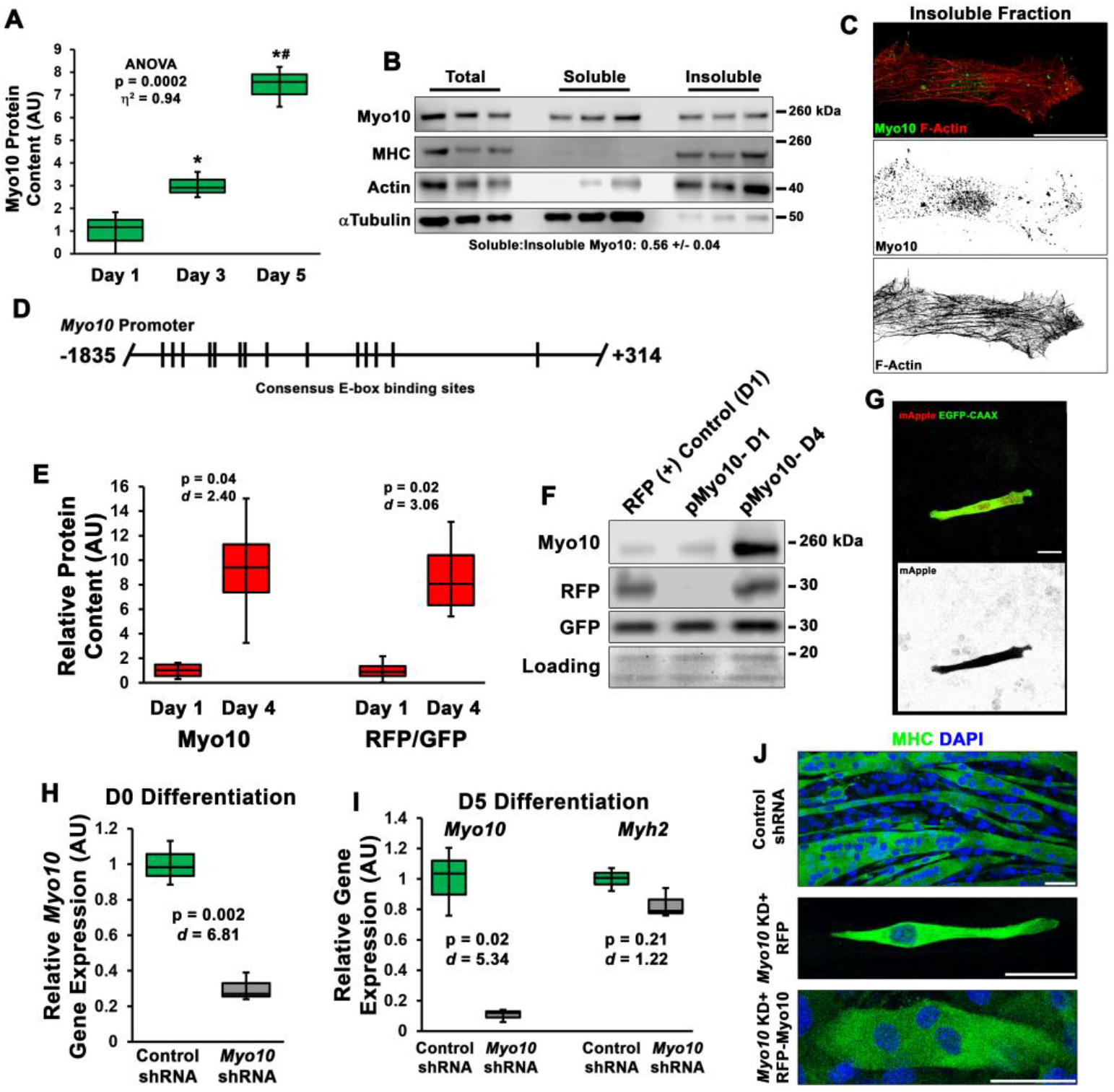
Myo10 is expressed by myoblasts during muscle differentiation and is required for myoblast fusion. (**A**) Myo10 protein content increases during myoblast differentiation, as measured by immunoblotting (n = 3 individual experiments). (**B**) Fractionation of differentiation day 5 myoblast cultures into soluble and insoluble cellular fractions reveals that the slight majority of Myo10 content exists in the soluble fraction. (**C**) Immunofluorescence (IF) of differentiated myoblast insoluble fractions shows that Myo10 of the insoluble cellular fraction is associated with the actin cytoskeleton (as shown by phalloidin staining) and can be found at the tips of thin cellular projections. (**D**) Schematic of the consensus E-box binding motifs (CANNTG) identified in the *Myo10* promoter. (**E-G**) Activation of the *Myo10* promoter reporter plasmid in differentiating myoblasts co-transfected with constitutively-expressed GFP-CAAX and mApple driven by the *Myo10* promoter depicted in **B** (n = 4 individual experiments). Efficient shRNA-mediated knockdown (KD) of myoblast *Myo10* gene expression in (**H**) undifferentiated and (**I**) differentiated myoblasts, whereas muscle differentiation is not affected by *Myo10* KD, as indicated by *Myh2* expression, a gene encoding a mature myosin heavy chain (MHC) expressed by skeletal muscle (n = 3 individual experiments). (**J**) Representative images of MHC IF of control shRNA cells, *Myo10* KD cells expressing a control RFP plasmid, and *Myo10* KD cells expressing a RFP-Myo10 rescue plasmid. Data are presented as box-and-whisker plots depicting 2^nd^ and 3^rd^ quartiles with minimum and maximum values. Data of **A** were analyzed using one-way ANOVA followed by Tukey post-hoc tests [α = 0.05; *p < 0.05 vs. Day 1 values; ^#^p < 0.05 vs. Day 3 values; effect size is presented as eta-squared (η^2^)]. Data of **E** and **H-I** were analyzed using two-tailed Welch’s T-tests with effect size presented as Cohen’s *d* (*d*). Scale bars represent (**G**) 10 or (**C, J**) 25 µm.

**Figure S3.**
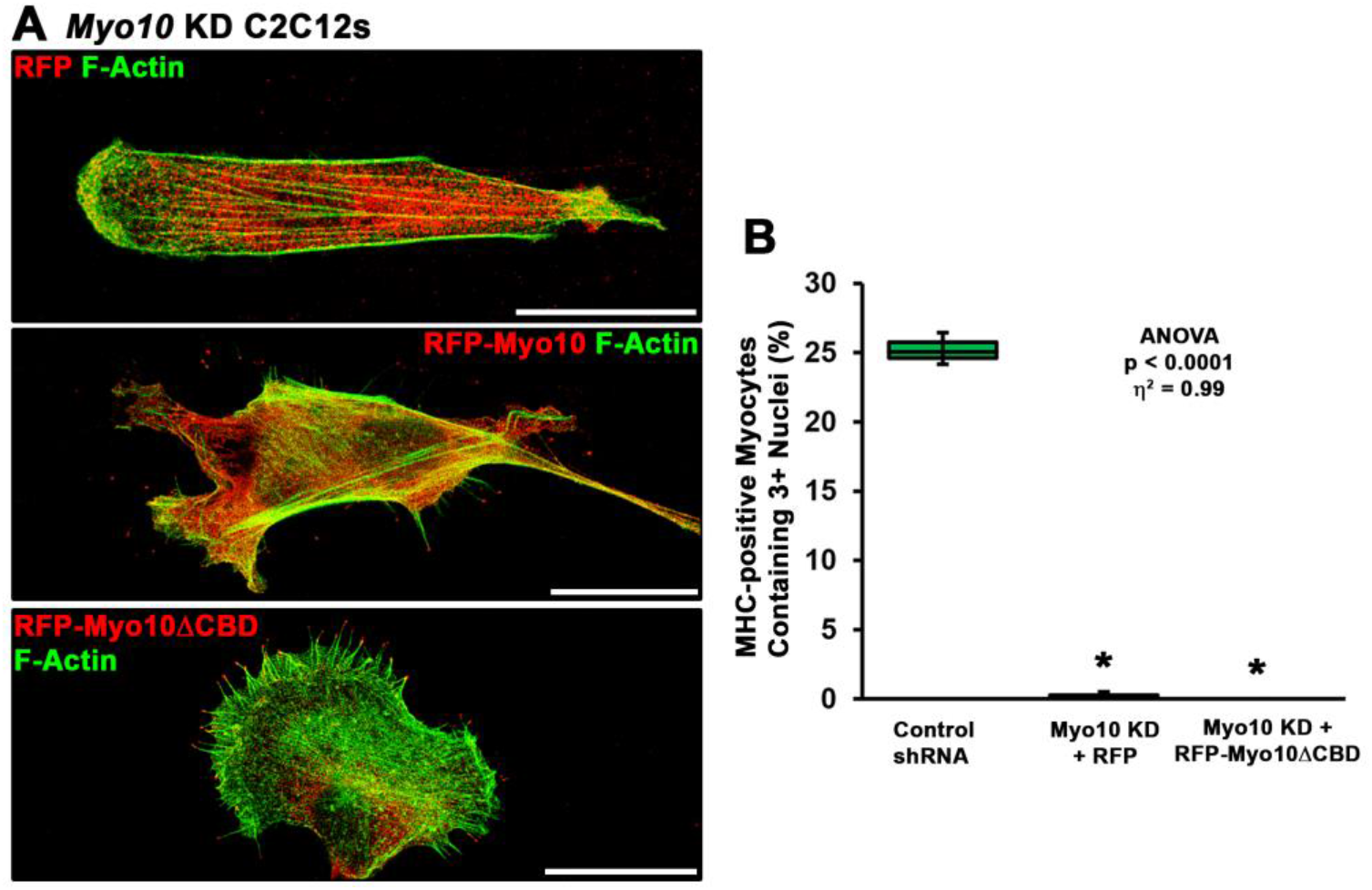
Rescue of myoblast fusion requires the Myo10 cargo binding domains. (**A**) Comparative images of *Myo10* KD myoblasts following transfection with RFP, RFP-Myo10, or a truncated RFP-Myo10 lacking C-terminal cargo binding domains (RFP-Myo10ΔCBD). Both RFP- Myo10 and RFP-Myo10ΔCBD expression result in filopodia formation. (**B**) Despite rescue of filopodia formation, RFP-Myo10ΔCBD does not rescue fusion ability of *Myo10* KD myoblasts (n = 6 independent experiments). Data were analyzed using one-way ANOVA followed by Tukey post-hoc tests [α = 0.05; *p < 0.05 vs. Control shRNA values; effect size is presented as eta-squared (η^2^)]. Scale bars represent 25 µm.

**Figure S4.**
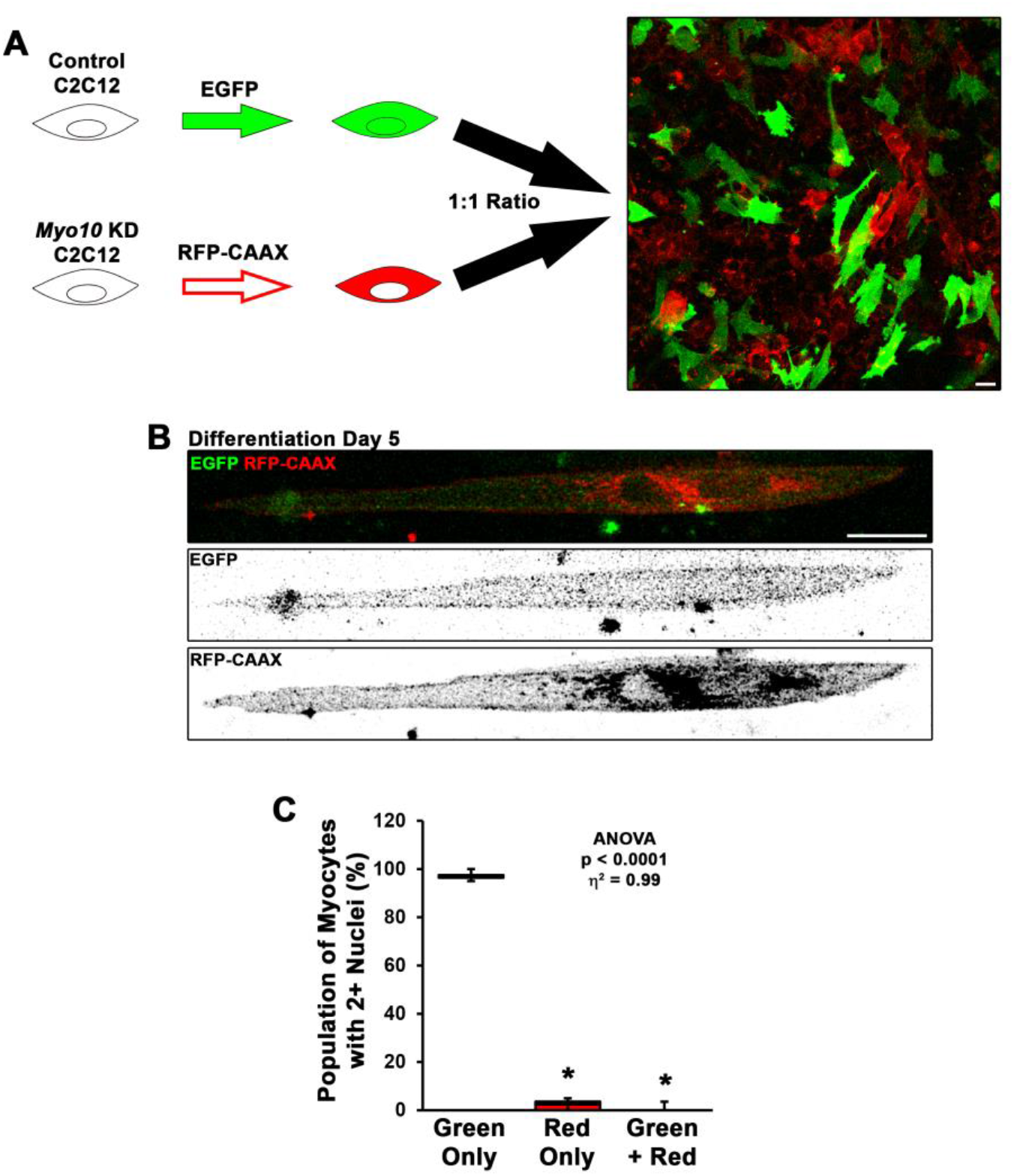
Fusion of myoblasts *in vitro* requires bilateral Myo10 expression. (**A**) To test if Myo10 is required unilaterally or bilaterally for myoblast fusion, control shRNA and *Myo10* KD C2C12 myoblasts were labeled with different variants of fluorescent proteins (EGFP and RFP-CAAX, respectively), then were mixed in a 1:1 ratio to assess prevalence of EGFP and RFP-CAAX incorporation into bi-/multi-nucleated myocytes (n = 5 independent experiments; bi- nucleated cells were included in this analysis because the fusion of *Myo10* KD cells with control cells will result in Myo10 KD in both nuclei). (**B-C**) Bi-nucleated cells having both EGFP and RFP-CAAX expression were exceeding rare in this assay, as only one instance was found in during the course of these experiments. Observed bi-nucleated *Myo10* KD cells having only RFP-CAAX expression are suspected to result from incomplete cytokinesis. Data were analyzed using one-way ANOVA followed by Tukey post-hoc tests [α = 0.05; *p < 0.05 vs. Green Only values; effect size is presented as eta-squared (η^2^)]. Scale bars represent 25 µm.

**Figure S5.**
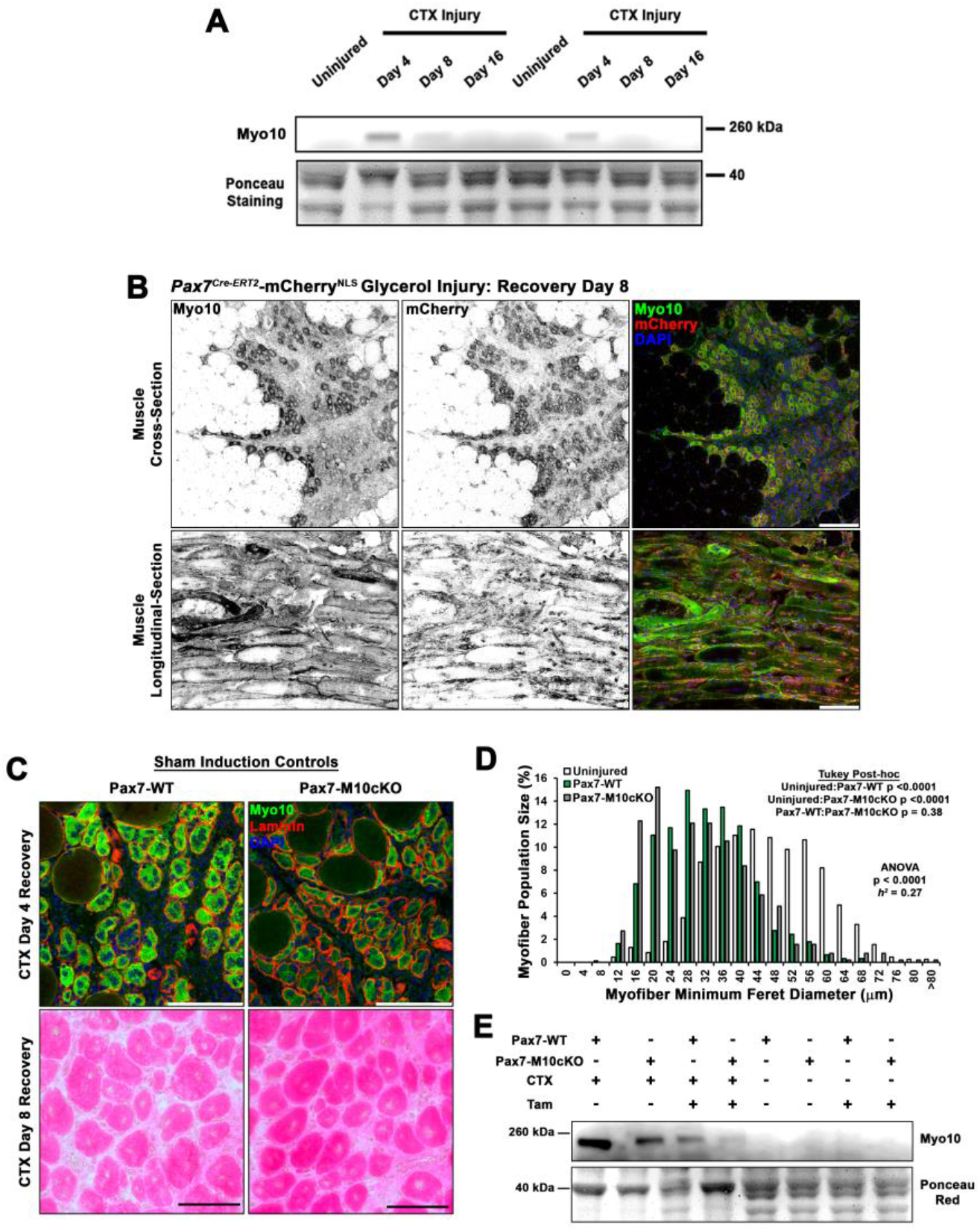
Myo10 is elevated during muscle regeneration. (**A**) Immunoblot for Myo10 content in uninjured and cardiotoxin (CTX)-affected tibialis anterior muscles after four, eight, and 16 days of regeneration. (**B**) Myo10 immunoreactivity is high in regenerating muscle of Pax7^Cre-ERT2^- mCherry^NLS^ mice 8 days following glycerol-induced injury. (**C-D**) Sham-induced Pax7^Cre-ERT2^ conditional *Myo10* knockout (Pax7-M10cKO; n = 4) mice and their non-floxed littermates (Pax7- WT; n = 5) do not demonstrate distinguishable Myo10 immunoreactivity or muscle regeneration following cardiotoxin (CTX)-induced injury of the tibialis anterior muscle. (**E)** Efficient ablation of full-length Myo10 protein in tamoxifen (Tam)-induced Pax7-M10cKO muscle following 4 days of regeneration from CTX injury achieved using the induction protocol depicted in Fig 4D. Data are displayed as a histogram of entire muscle fiber populations and were analyzed using one-way ANOVA followed by Tukey post-hoc tests (α = 0.05). Scale bars represent 100 µm.

**Figure S6.**
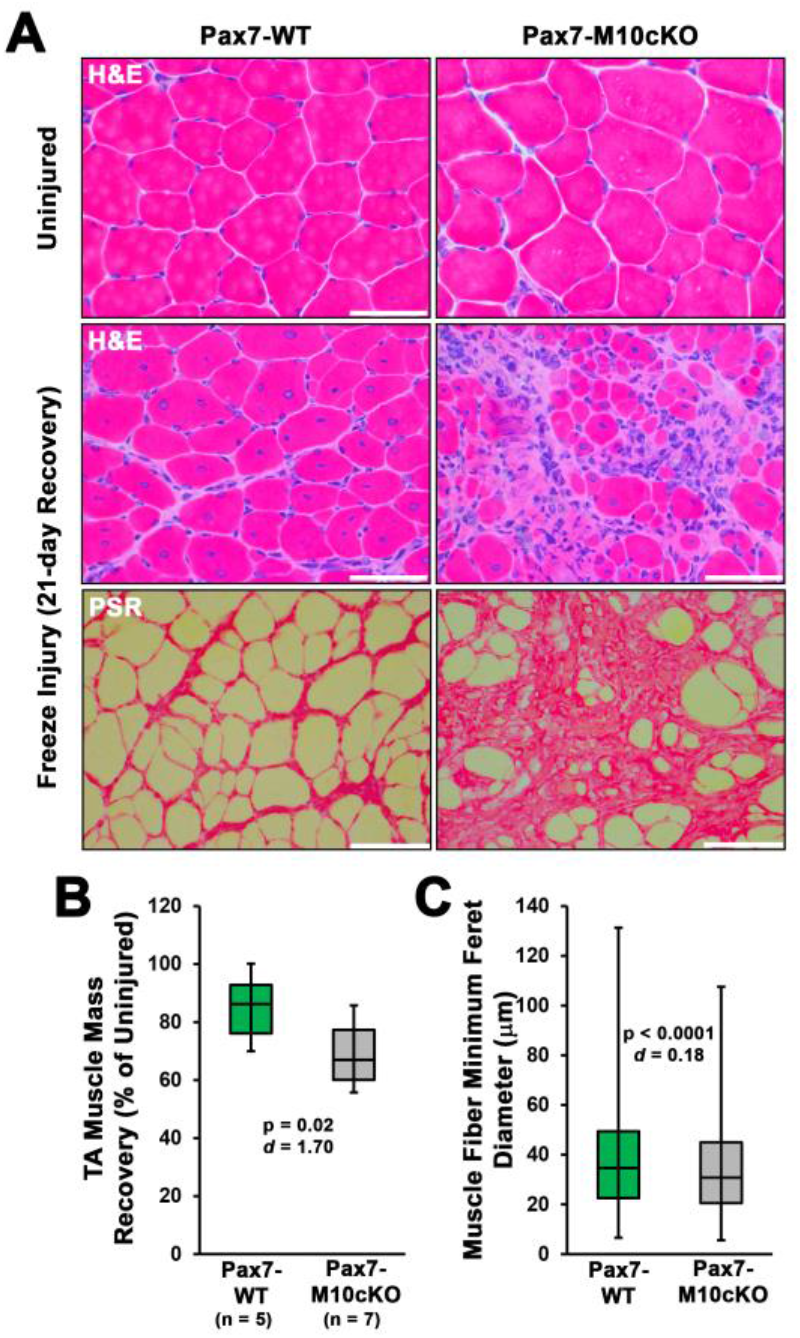
Satellite cell ablation of Myo10 impairs muscle regeneration from freeze injury. Pax7-WT and Pax7-M10cKO littermate mice were induced with tamoxifen, and subjected to freeze injury of the tibialis anterior (TA) muscle (n = 5-8). (**A**) Following 21 days of recovery, Pax7- M10cKO (n = 8) mice exhibit impaired regeneration, as shown by H&E staining, and increased fibrosis, as shown by picrosirius red (PSR) staining. (**B**) Muscle mass and (**C**) muscle fiber size recovery was diminished by loss of satellite cell Myo10 (n = 12065-16707 fibers). Data are presented as box-and-whisker plots depicting 2^nd^ and 3^rd^ quartiles with minimum and maximum values, and are analyzed using two-tailed Welch’s T-tests with effect size presented as Cohen’s *d* (*d*). Scale bars represent 100 µm.

**Figure S7.**
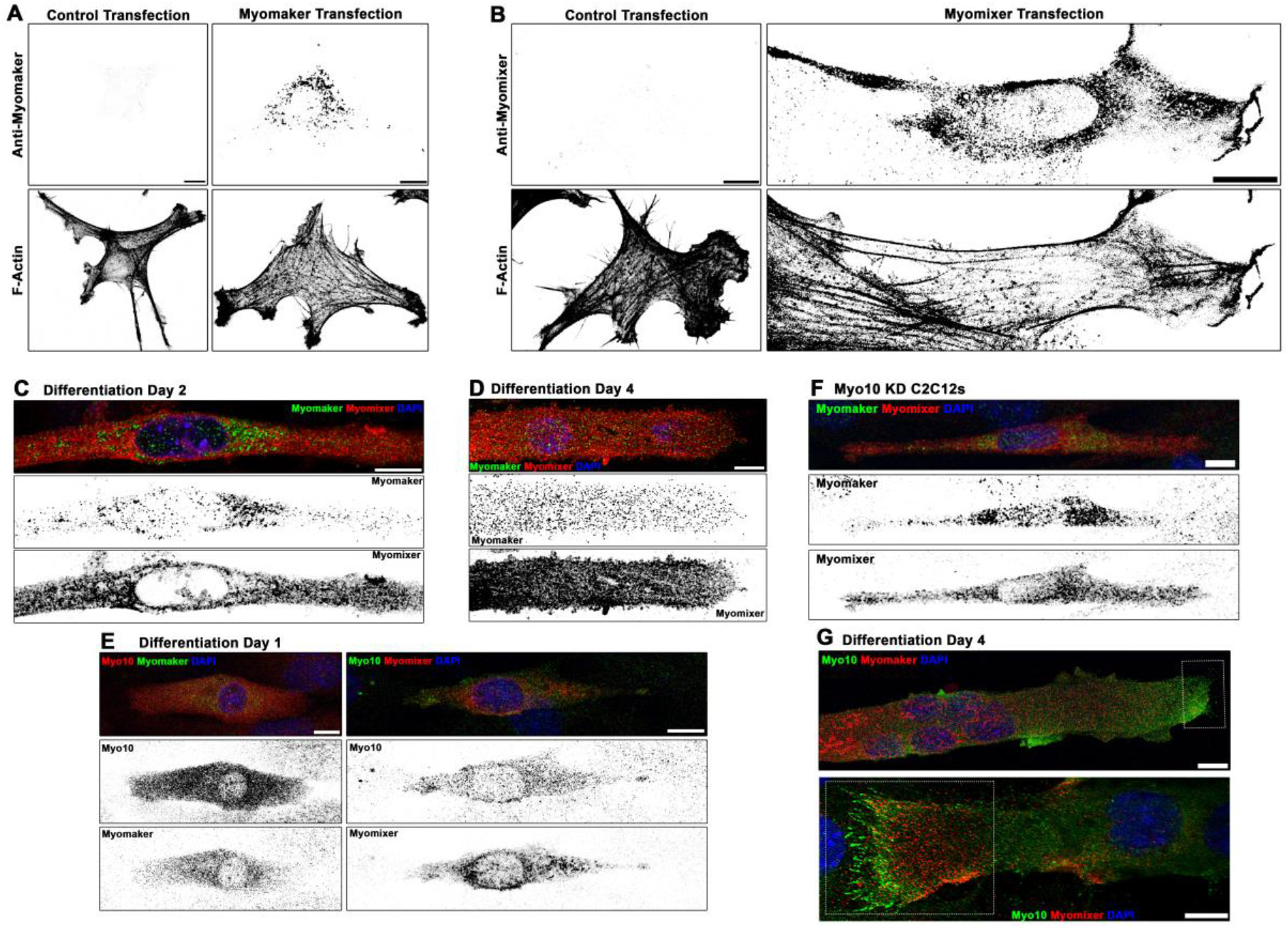
Localization patterns of Myomaker and Myomixer in differentiating myoblasts. Specificity testing of commercially-available (**A**) Myomaker and (**B**) Myomixer antibodies for immunofluorescent detection of these proteins following transfection of undifferentiated myoblasts. IF staining pattern of Myomaker and Myomixer (**C**) early in myogenic differentiation and (**D**) in fully differentiated myotubes. Myomixer antibody was applied prior to cell permeabilization. (**E**) At day 1 of differentiation, myoblasts co-expressing Myo10 with Myomaker and Myomixer are observed. (**F**) Differentiated *Myo10* KD cells express both Myomaker and Myomixer. (**G**) Localization patterns of Myo10 with Myomaker (top) and Myomixer (bottom) in differentiated myotubes (insets are found in Figure 5B). Scale bars represent 10 µm.

**Figure S8.**
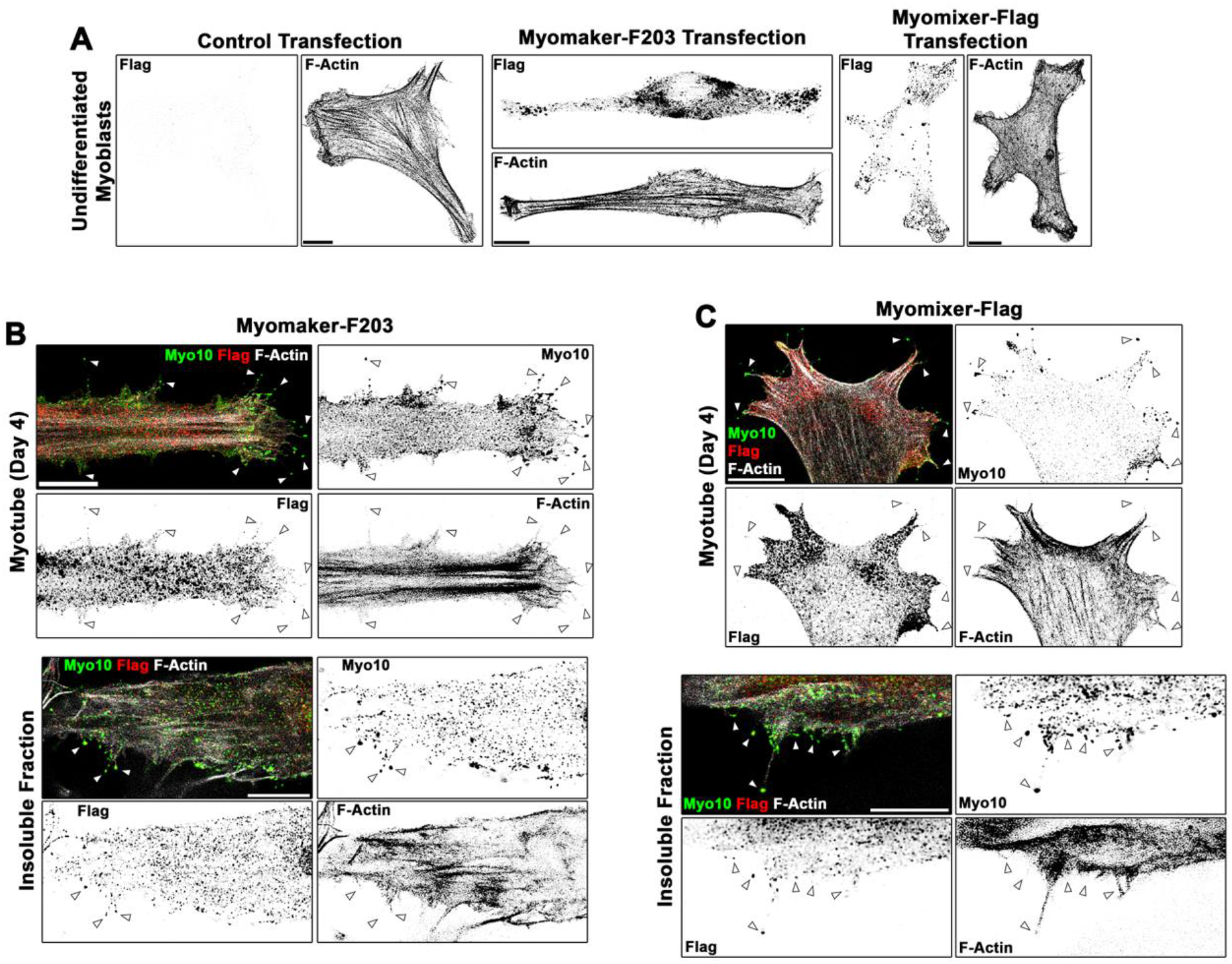
Localization patterns of Flag-tagged Myomaker and Myomixer constructs in differentiating myoblasts. (**A**) Anti-Flag staining of undifferentiated myoblasts following transfection of control, Myomaker-F203, or Myomixer-Flag plasmids reveals specificity of the anti-Flag antibody and that these Flag-tagged versions of Myomaker and Myomixer localize similarly to their wild-type counterparts (shown in **Figure S7**). Investigation of the localization of (**B**) Myomaker-F203 and (**C**) Myomixer-Flag in differentiated myotubes reveals Flag-positive puncta observed in Myo10-positive filopodia of both whole cells and the insoluble cellular fraction. Scale bars represent 10 µm.

## SUPPLEMENTAL MOVIE CAPTIONS

**Supplemental Movie 1. Movement of undifferentiated myoblasts.** Time-lapse confocal imaging of undifferentiated myoblasts expressing RFP-CAAX. Individual frames were utilized to make **Figure 1A**. Images were acquired every 15 minutes. Scale bar represents 25 µm.

**Supplemental Movie 2. Cellular projections of undifferentiated myoblasts.** Representative time-lapse movies of cellular projections from the anterior, dorsal, and posterior positions of undifferentiated myoblasts expressing RFP-CAAX. Individual frames are included in **Figure S1A**. Images were acquired every 20 seconds for a duration of 2 minutes.

**Supplemental Movie 3. Morphology changes of differentiating myoblasts.** Time-lapse movie of differentiating myoblasts expressing RFP-CAAX. Images were acquired every 15 minutes. Scale bar represents 100 µm.

**Supplemental Movie 4. Myotube lateral edge projections.** Time-lapse video of the lateral edge of a differentiated myotube expressing RFP-CAAX. Individual frames are included in **Figure 1B**. Images were acquired every minute.

**Supplemental Movie 5. Non-fluorescent detection of myotube lateral edge projections.** Time-lapse of differential interference contrast imaging showing dynamic cellular projections at the lateral edge of a myotube. The arrow indicates where pronounced projections are clearly visible. Images of this myotube from later time points are also found in **Figure 1D** and **Supplemental Movie 8**. Images were acquired every 30 seconds.

**Supplemental Movie 6. Dynamic myotube dorsal protrusions.** Time-lapse confocal imaging of a GFP-CAAX-expressing myotube exhibiting dynamic dorsal protrusions. Individual frames are included in **Figure 1B.** Images were acquired every 30 seconds. Scale bar represents 25 µm.

**Supplemental Movie 7. Dynamics of myotube cellular projections.** Time-lapse confocal imaging of diverse cellular projection of differentiating myotubes expressing GFP-CAAX. Individual frames were utilized to make **Figure 1A**. Images were acquired every 20 minutes.

**Supplemental Movie 8. Cellular fusion at the myotube lateral edge.** Time-lapse confocal images of a GFP-CAAX-expressing myotube initiating fusion with fine protrusions extending from the lateral edge. Individual frames are included in **Figure 1F**. Images were acquired every 15 minutes. Scale bar represents 25 µm.

**Supplemental Movie 9. Myotube fusion initiated by a lamellipodial extension.** Time-lapse confocal images of a GFP-CAAX-expressing myotube initiating fusion using a lamellipodial extension adorned with fine protrusions. Individual frames are included in **Figure 1E**. Images were acquired every 15 minutes. Scale bar represents 25 µm.

**Supplemental Movie 10. Cellular fusion initiated by a lamellipodial extension.** Time-lapse confocal images of a GFP-CAAX-expressing myotube initiating fusion using a lamellipodial extension. Images were acquired every 15 minutes.

**Supplemental Movie 11. Activation of the *Myo10* promoter in differentiating myoblasts.** Time-lapse confocal images of a myoblast co-expressing GFP-CAAX and a reporter plasmid consisting of mApple driven by the *Myo10* promoter at day 1 of differentiation. Individual frames are included in **Figure S2G**. Images were acquired every 20 minutes. Scale bar represents 10 µm.

**Supplemental Movie 12. Loss of Myo10 in myocytes prevents filopodia formation.** Confocal images of cellular projections exhibited by differentiated control and *Myo10* shRNA knockdown myocytes. Individual frames are included in **Figure 3D**. Images were acquired every 10 seconds. Scale bar represents 5 µm.

**Supplemental Movie 13. Loss of Myo10 does not prevent lamellipodial extension formation.** Differentiating Myo10 knockdown myoblasts expressing RFP-CAAX produce lamellipodial extensions during differentiation. Images were acquired every 20 minutes. Scale bar represents 10 µm.

